# Molecular control of interfacial protein structure on graphene-based substrates steers cell fate

**DOI:** 10.1101/2020.02.11.944678

**Authors:** Sachin Kumar, Sapun H. Parekh

**Affiliations:** Department of Biomedical Engineering, University of Texas at Austin, 107 W. Dean Keeton Rd., Austin, TX 78712, USA; Department of Molecular Spectroscopy, Max Planck Institute for Polymer Research, Ackermannweg 10, D-55128 Mainz, DE

**Keywords:** Graphene-based materials (GBMs), protein interaction, protein molecular structure, protein bioactivity, stem cell differentiation

## Abstract

The use of graphene-based materials (GBMs) for tissue-engineering applications is growing exponentially due to the seemingly endless multi-functional and tunable physicochemical properties of graphene, which can be exploited to influence cellular behaviours. Despite many demonstrations wherein cell physiology can be modulated on GBMs, a clear mechanism connecting the different physicochemical properties of different GBMs to cell fate has remained elusive. In this work, we demonstrate how different GBMs can be used to cell fate in a multi-scale study – starting from serum protein (Fibronectin) adsorption to molecular scale morphology, structure and bioactivity, and finally ending with stem cell response. By changing the surface chemistry of graphene substrates with only heating, we show that molecular conformation and morphology of surface adsorbed fibronectin controls epitope presentation, integrin binding, and stem cell attachment. Moreover, this subtle change in protein structure is found to drive increased bone differentiation of cells, suggesting that physicochemical properties of graphene substrates exert cell control by influencing adsorbed protein structure.

## 1. Introduction

Graphene-based substrates are emerging as promising next generation biomaterials for tissue engineering applications. Many studies have shown promising results where different cells respond favorably to graphene-based materials (GBM), demonstrating strong biocompatibility.^1-3^ Particularly for tissue engineering and regenerative medicine, stem cell differentiation has been reported to change in response to different GBMs.^4^ Researchers have used GBMs in various forms: as films, in porous scaffolds, as substrate coatings, and in reinforced materials for stem cell based tissue engineering applications.^2,3^ While many of these studies highlight the significant impact of graphene nanoparticles and surfaces on stem cell function, a clear mechanism connecting the different physicochemical properties of GBMs with stem cell fate is yet unknown. Nevertheless, studies have shown that roughness, surface chemistry, wettability, conductivity and mechanical properties of graphene surfaces all modulate cell fate,^2,5,6^ but how each of these properties uniquely influence stem cell is not well understood. Given the large enumeration of results in this area, a strong need exists to better relate graphene substrate physicochemistry with cell behaviour from a mechanistic picture to highlight the essential features of GBMs that can tune cell fate.

It is well documented that nearly all biomaterials adsorb proteins upon contact with blood or serum. The molecular adsorption of proteins and other molecules to biomaterials can strongly control how cells perceive and react to biomaterials. GBMs are no different and have been shown to adsorb numerous proteins on their surfaces, similar to other biomaterials, which no-doubt modify the GBM ultimately presented to the cells.^7-10^ Chong *et al.* demonstrated that, depending on surface properties of graphene oxide (GO) and reduced graphene oxide (RGO), the serum proteins bovine serum albumin (BSA), fibrinogen (FG), Transferrin (Tf) and Immunoglobulin (Ig) not only adsorbed, but also interacted and oriented differently.^11^ The interaction and orientation of adsorbed proteins on the GBM substrates then presented the biochemical and topographical interface for stem cell interaction. The sensitive nature of stem cells to the interfacial protein structural matrix via cell surface receptors^12^ (such as integrins) makes elucidating and controlling the interaction and conformation of ECM proteins with GBMs critical for deterministically controlling cell fate response. Hence, a complete picture showing how GBMs with different physicochemistry modify ECM protein interaction and conformation and downstream stem cell function is necessary.

Despite its importance, surprisingly few studies have highlighted the role of adsorbed interfacial protein on graphene surfaces influencing stem cell response.^13-15^ Lee *et al*. demonstrated that adsorbed insulin on pristine graphene undergoes denaturation whereas insulin on GO retained its conformation and activity. As a result, the authors suggested that GO provided active pre-concentrated insulin for stem cell interaction, resulting in better adipogenic differentiation.^13^ In addition, the authors used computer simulation and circular dichroism (CD) spectra of insulin extracted from the GBM surfaces to comment on insulin structure. From this study, it is tempting to link the substrate-induced structure of insulin to more adipogenic differentiation; however, no identification of biochemical activity changes of insulin on graphene and GO surfaces was presented. Experiments directly probing protein molecular structure, conformation, and exposure of bioactive domains of proteins are required to unequivocally demonstrate if GBMs modulate protein structure to influence stem cell interaction. In addition, controlled experiments evaluating the effect of the adsorbed protein and its molecular features on stem cell attachment and differentiation should be done in absence and presence of serum. Providing such experiments will provide mechanistic insight on GBM-mediated stem cell response for future rational design of materials from graphene.

Here, we study ECM protein adsorption, interaction, orientation and structure from the macroscale to the molecular scale on GBMs and perform stem cell culture in the presence and absence of serum to relate the physicochemistry of GBMs with protein structure and cell fate. We used GO and RGO – the two most commonly used GBMs as exemplary engineering materials – to evaluate bioactivity of GBMs having different surface chemistries by investigating the effect of GBM chemistry on fibronectin (FN – a protein abundant in the ECM), adsorption, orientation, and structure. We couple these surface characterization experiments with biochemical activity assays and stem cell response experiments, allowing us to elucidate the link between physicochemical properties of GBMs, FN protein unfolding, altered biochemical activity, and stem cell differentiation.

## 2. Results and discussion

### 2.1. Preparation and characterization of GBM substrates

GO and RGO coated glass substrates were prepared by spin coating GO on to cleaned glass substrates (**Figure S1**). Spin coating has been reported to provide a uniform film thickness of GO over a large area with high reproducibility^16^ due to rapid evaporation of solvent that improves interaction between GO and hydrophilic substrates. This process provides strongly adhered, continuous films with minimal wrinkling.^16,17^ We produced continuous GO films having a thickness of ∼ 5.5 nm on UVO-treated glass (**Figure S2(a)**). RGO was produced from GO substrates by classical low thermal reduction of GO substrates^18,19^ (Explained in detailed in Supplementary Information). RGO substrates had a blackish appearance (compared to a slight brown tint of GO), consistent with reduction of oxide groups (**Figure S1**) and verified with XPS (**Figure 1(a)** and **Table 1**).

**Table 1:** Quantification of different chemical functional groups present on GO and RGO surface

|  | GO (%) | RGO (%) |
| --- | --- | --- |
| C=C/C-C | 54.5 | 71.0 |
| C-OH | 11.9 | 13.1 |
| C-O-C | 16.5 | 7.1 |
| -C=O and HO-C=O | 17.1 | 8.8 |

**Figure 1:**
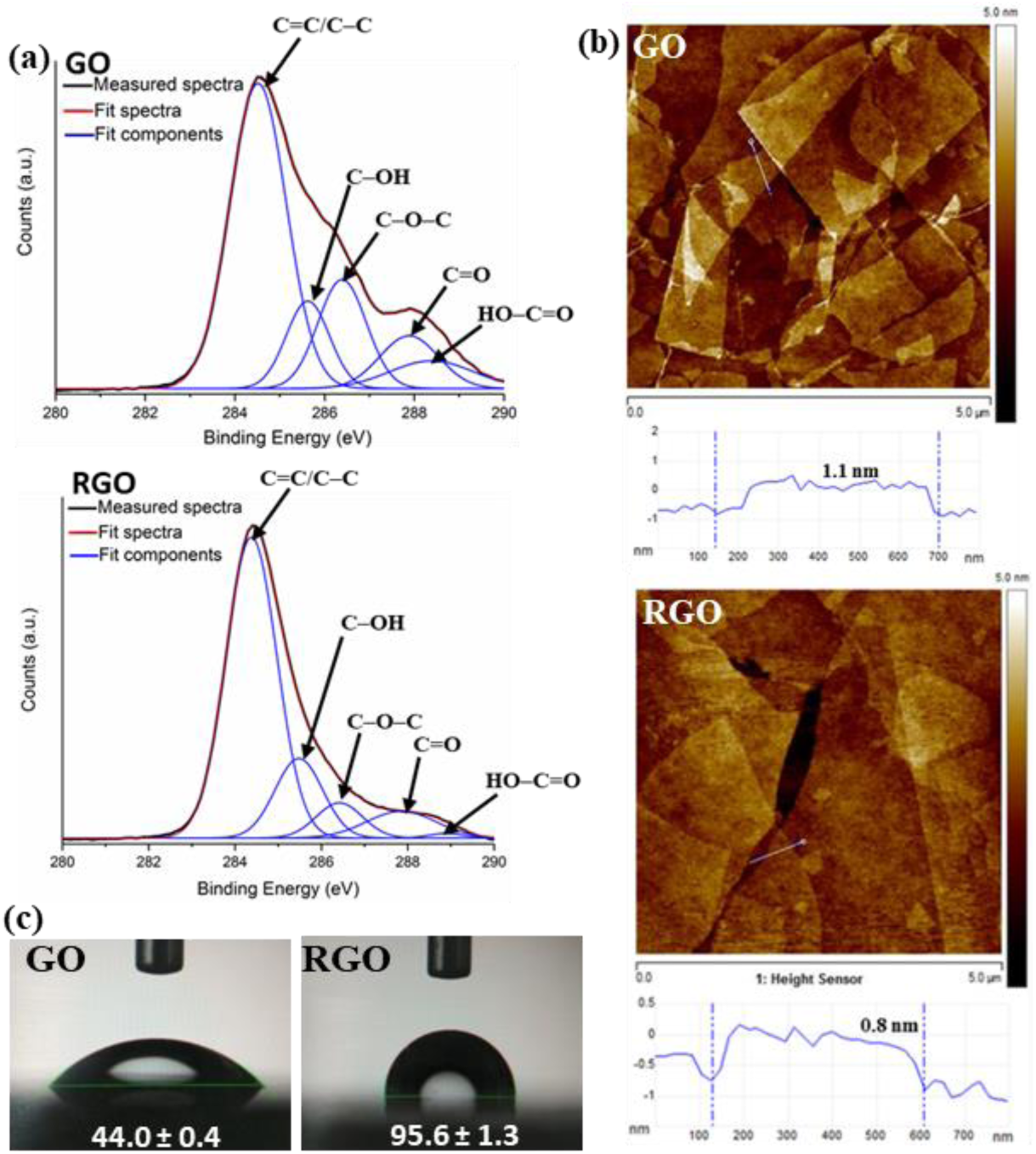
Physical and chemical characterization of GBMs. (a) Deconvolution of XPS C1s core-level spectra of GO and RGO into various chemical functional states using a Gaussian fitting after base line correction. (b) AFM images show surface morphology of GO and RGO flakes on glass along with height profile. (c) Static water contact angle micrograph on GO and RGO surface shows clear differences in hydrophilicity.

Physicochemical characterization of GO and RGO substrates showed coated flakes were smooth and continuous with distinguishable edges of individual sheets lying flat on a glass surface without wrinkles (**Figure S2 (b) and (c)**). Importantly we found that thermal reduction of GO to RGO did not produce any significant morphological difference apart from apparent decrease in flakes thickness (**Figure 1(b)**), which is consistent with previous work showing that the thickness of an individual GO sheet reduced from 1.1 nm to 0.8 nm after thermal reduction.^20,21^ Upon thermal reduction more than 50% of epoxy, carbonyl and carboxyl functional groups were removed; however, the hydroxyl functional groups were almost unchanged (**Figure 1(b) and Table 1**). Studies have shown that highly stable hydroxyls and carbonyl groups are difficult to remove from the graphene sheet edges at low temperature reduction without inducing detects;^18,19^ as a result, RGO retains 15 - 25 % residual oxygen concentration upon low thermal reduction. Additional characterization with Raman spectroscopy of the intensity ratio of D and G bands (I_D_/I_G_) of GO and RGO showed that removal of oxygenated functional groups partially restored defects in GO (**Figure S2 (e)**). From a functional perspective, thermal reduction resulted in an increase hydrophobicity indicated by a contact angle of 95.6 ± 1.3 for RGO in comparison to 44.0 ± 0.4 for GO (**Figure 1(c)**). We note that during reduction only a small difference in surface roughness was observed between GO and RGO (Supplementary Information). Hence, it was reasonable to assume the difference in contact angle was mainly due to a change in surface chemistry. Overall, thermal reduction of GO to RGO removed oxygenated functional groups and resulted in two different GBMs, having similar nearly identical physical properties (roughness and defects) but different chemical surface properties (hydrophilicity).

### 2.2. Fibronectin protein adsorption on GO and RGO coated substrates

Real-time FN adsorption on GO and RGO substrates was studied using QCM-D. **Figure 2(a)** shows the frequency shift (decrease) with time in response to adsorption of FN on GO and RGO substrates; the frequency shift is linearly correlated with the mass of material adsorbed on the QCM sensor to first order. FN adsorption on GO and RGO substrates showed different kinetics and overall adsorption: a faster change in frequency was observed on RGO substrate in comparison to GO, and the GO and RGO substrates reached final frequency saturation at −60 and −298 Hz, respectively, corresponding to mass density of 354 ng/cm^2^ and 1758 ng/cm^2^ of adsorbed FN. Clearly. RGO showed a more significant amount of adsorption of FN (nearly 5 times), which is in good agreement with the data obtained from the XPS results (**Figure S3(a)**). The effective thickness of adsorbed FN layer on GO and RGO substrates was calculated using the Voight model reported in literature for FN adsorption.^22-24^ Based on this model, the adsorbed FN formed an effective layer of thickness of 3.7 nm on GO in comparison to 15.2 nm on RGO. In order to confirm the number from the model, AFM images of FN adsorbed on GO and RGO substrates (**Figure 2(b))** were obtained. AFM micrographs show flaky features from neat GO and RGO substrates, and FN coated GO showed partial and thinner FN coverage (max thickness ∼ 5 nm) in comparison to the RGO that showed denser and thicker FN coverage (max thickness ∼ 16 nm). These results confirm that FN forms a denser, multilayer coating on RGO compared to GO.

**Figure 2:**
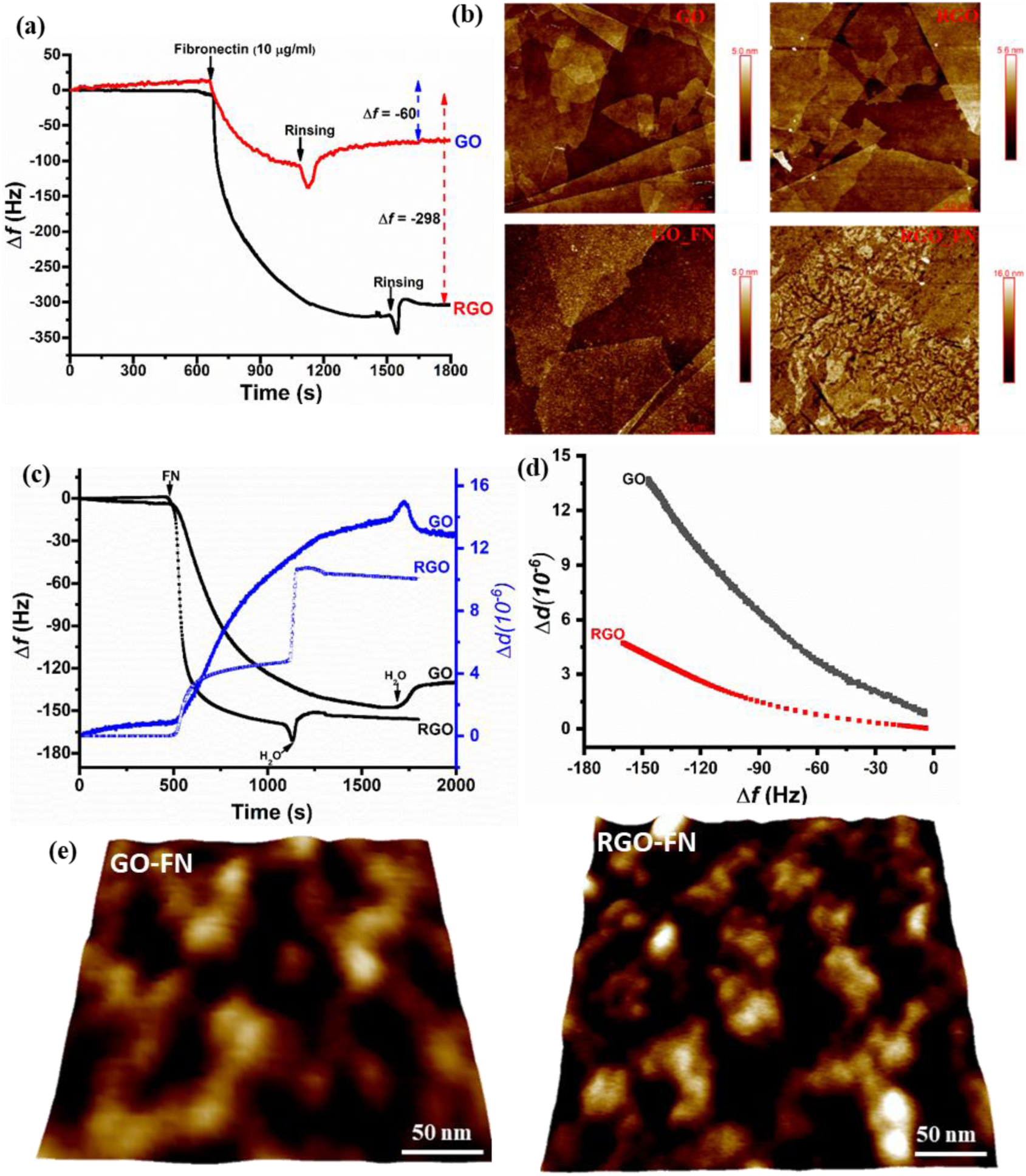
FN adsorption and optimization for homogeneous coating of FN on GBM substrates. (a) Real-time QCM adsorption profile of FN. (b) AFM micrograph of before and after FN adsorbed GO and RGO substrate. (c) QCM-D profile of optimized FN adsorption on GO (20 µg/ml) and RGO (5 µg/ml) surface. (d) QCM dissipation per frequency (ΔD/ΔF) plot of FN on GO and RGO surface. (e) High resolution AFM micrograph recoding of structural morphology of FN on GO and RGO surface.

Higher adsorption of FN on RGO can be attributed to the hydrophobic nature of RGO. Hydrophobic surfaces having lower surface energy mediated significantly more FN adsorption in comparison to hydrophilic surfaces with high surface energy.^25^ This is believed to occur because of energetically favorable displacement of water molecules on graphene hydrophobic surfaces (having low surface energy) by proteins.^26,27^ In addition, FN (having an isoelectric point at pH 5.92) is negatively charged at physiological pH (7.4), which favors more electrostatic adsorption on surfaces with less negative surface charge.^28^ GO, having a rich amount of oxygenated functional groups, could provide more negative charge surface at physiological pH in comparison to RGO, resulting in less attachment to GO.

The heterogeneous and differential amount of protein adsorption can complicate the interpretation of bioactivity and biological studies, so we optimized the solution concentration of FN to obtain a more homogeneous and similar amount of FN adsorption on both GBMs. We found that a FN solution concentration of 20 µg/ml produced for GO adsorption resulted in similar FN mass density (692 ng/cm^2^) as a 5 µg/ml FN solution on RGO (805 ng/cm^2^) (**Figure 2(c)**). These surfaces showed uniform thickness by AFM with 5.6 nm and 7.4 nm for GO and RGO, respectively (**Figure S3 (b)**).

In addition to adsorbed mass density and thickness, QCM-D allows one to study the adsorption energy by quantifying the dissipation per frequency (ΔD/ΔF). The (ΔD/ΔF) quantity is related to hydrodynamic size, packing, conformation, and strength of surface-bound molecules, with larger (ΔD/ΔF) corresponding to more “floppy” and weakly attached species.^29-31^ **Figure 2(d)** shows a plot of dissipation changes per frequency changes (ΔD/ΔF) for GO and RGO upon FN adsorption. GO surfaces showed a lower adsorption rate during FN adsorption (**Figure 2(c)**) with higher (ΔD/ΔF) (**Figure 2(d)**), which suggests that FN molecules on GO are more loosely packed and weakly adsorbed compared to those on RGO.^32^ As described above, the hydrophilic GO surface having more negative surface charge can provide enhanced surface hydration leading to fewer FN adhesion sites, allowing for a more flexible FN structure on the surface. On the other hand, RGO surface showed lower (ΔD/ΔF) and faster FN adsorption, indicating a more rigid and compact structure during adsorption. We used AFM to further measure the size and morphology of FN on GO and RGO surfaces at high resolution (**Figure 2(e))**. Interestingly, FN on GO showed elongated fibrillar structures with an apparent formation of a protein network, whereas FN on RGO appeared as isolated and compact in globular structures. Analyzing the characteristic length of the FN structures GO and RGO (n∼80 on each surface), we found a length (L_FN_) of 108.2 ± 28.1 nm (in agreement with expected length of Fn molecule with elongated morphology nm ^33^) and L_FN_ of 53.1 ± 18.6 nm, respectively. The smaller L_FN_ on RGO is consistent with a stronger, more packed molecular structure inferred from QCM-D measurements.

To verify our hypothesis that FN molecular structure was elongated on GO and compact on RGO, we used nano-IR AFM (nAFM-IR), a technique that can provide nanoscale IR spectroscopic information. IR spectroscopy has been well established to report on protein secondary structure^34^ and combined with AFM, we report the first measurements of FN secondary structure on a surface in **Figure 3(a)**. The AFM topographic and nAFM-IR absorbance image at 1550 cm^−1^ for FN on GO and RGO are shown for comparison.

**Figure 3:**
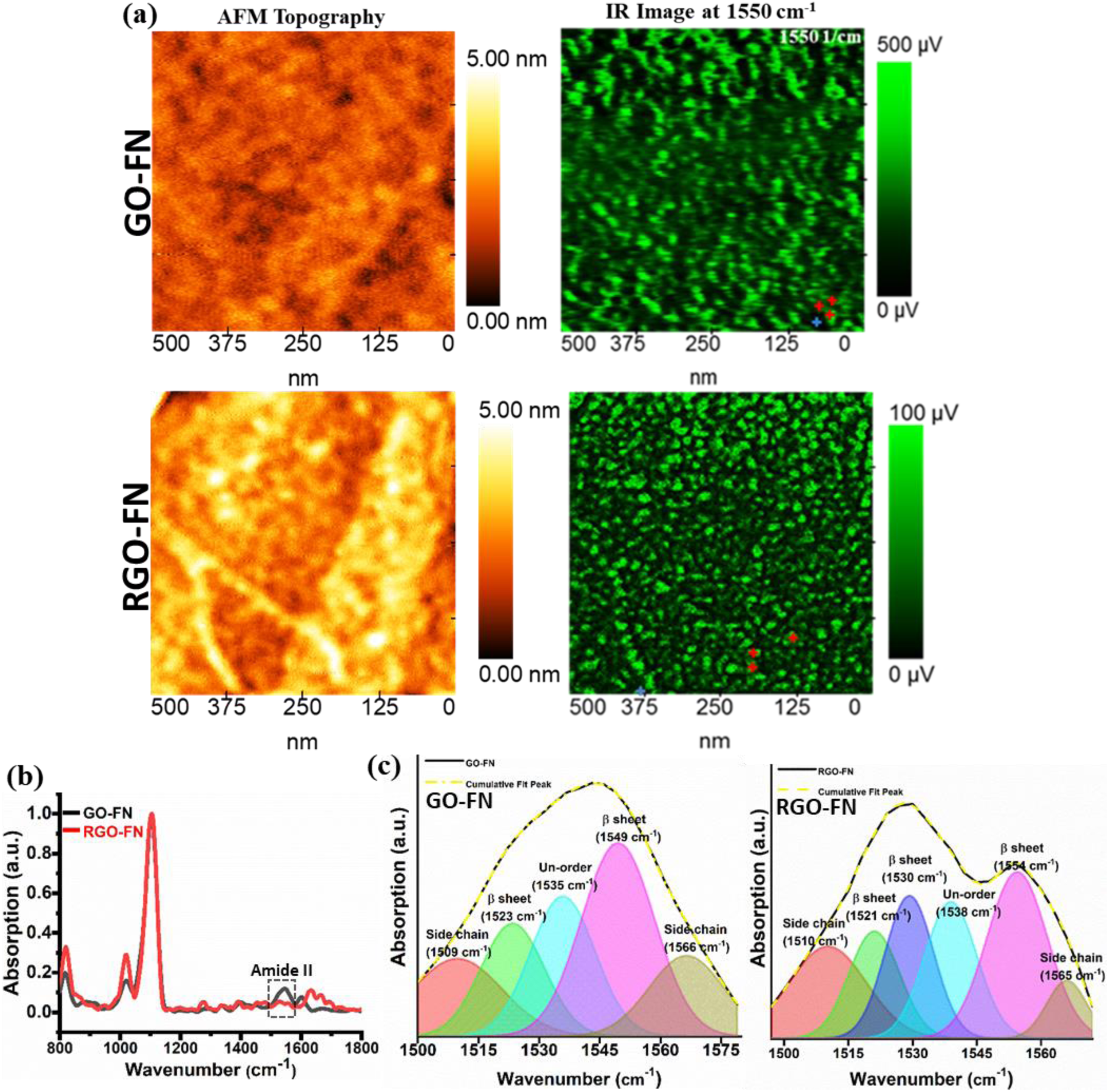
nAFM-IR shows molecular and morphological changes of adsorbed FN on GBMs. (a) nAFM-IR topography and IR chemical profile of FN on GO and RGO surface. (b) Single spot IR spectra of FN adsorbed on GBMs; each spectra is an average of three red spots (indicated in IR image at 1550 cm^−1^). (c) spectral deconvolution of amide II band of FN on GO and RGO.

The nAFM-IR image at 1550 cm^−1^ clearly shows an elongated fibrillar structure of FN on GO surface whereas FN on RGO shows a compact, round structure. This result is consistent with the high resolution AFM in **Figure 2(e)**. In addition, we obtained IR spectra from individual FN features on the GO and RGO surface, which showed clear differences in the Amide II band, suggesting a molecular structural change of the protein on the two GBMs (**Figure 3(b)**). FN on GO showed a broad single peak centered at 1545 cm^−1^ corresponding to a disordered β-sheet. On the other hand FN on RGO showed doublet amide II peak with maxima at 1528 and 1556 cm^−1^, corresponding to two β-sheet peaks.^35^ We note that FN is natively a β-sheet rich protein, showing ∼79% β-sheet ^36^. Decomposition of the Amide II spectra (**Figure 3(c) and Table 2**) quantitatively showed that FN adsorbed on GO was composed of 49.1% β sheet, 20.8% unordered and 30.1% turn, having β sheet to unordered ratio of 2.3. On the other hand, FN on RGO surface showed 57.7% β sheet, 18.4% unordered and 33.9% turn having β sheet to unordered ratio of 3.1. Thus on GO, the β sheet amount decreased by 26% and the resulting increased unordered structure demonstrates that the FN adsorbed to GO was unfolded compared to that on RGO surface. Taken together, the above results the show how the small change in chemistry (mainly different fraction of oxygenated functionalities) on GO and RGO cause adsorbed FN to have distinct size, conformation, and molecular structure. Hence, different physicochemical properties of GBMs can cause adsorbed proteins to have different interaction, morphology, conformation, and molecular structure.

**Table 2:** Secondary structural components of adsorbed FN on GO and RGO surface

|  | GO-FN (%) | RGO-FN (%) |
| --- | --- | --- |
| β-sheet | 49.1 | 57.7 |
| Un-order | 20.8 | 18.4 |
| Turn | 30.1 | 23.9 |

### 2.3. Bioactivity of fibronectin coated GO and RGO

With FN adsorbed to GO and RGO showing distinct morphology and molecular structure, we next evaluated the bioactivity of the adsorbed protein, which is known to depend on the structural conformation of the protein.^37^ FN is an ECM protein that is known to possess cryptic, or normally hidden, cysteines as well as other functional binding domains (such as integrin binding sites). Hence, we evaluated if adsorption of FN to GO and RGO exposed bioactive binding sites starting with cryptic cysteine residues. We used maleimide-conjugated Alexa-488 (MA) that binds specifically with free sulfhydryl groups to determine if adsorption of FN molecules to either GBM exposed any cryptic cysteines.

FN has two nearly identical subunits, each possessing two cryptic cysteine units at FnIII7 and FnIII15 (as illustrated in **Figure 4(a)**).^38^ **Figure 4(b)** shows representative images of MA staining on FN molecules on GO and RGO surfaces. FN on GO surface showed significantly more fluorescence MA (green color) staining in comparison to RGO surface. Quantifying the fraction of image areas stained by MA, GO showed 78% labeling of FN by MA whereas FN on RGO only 20% of the available labeled – even though the protein density on the two surfaces was nearly identical. This result clearly indicates more accessibility of cryptic cysteine groups of FN on GO, which is in agreement with the elongated fibril structure of FN on GO that was more unfolded compared to compact FN on RGO.

**Figure 4:**
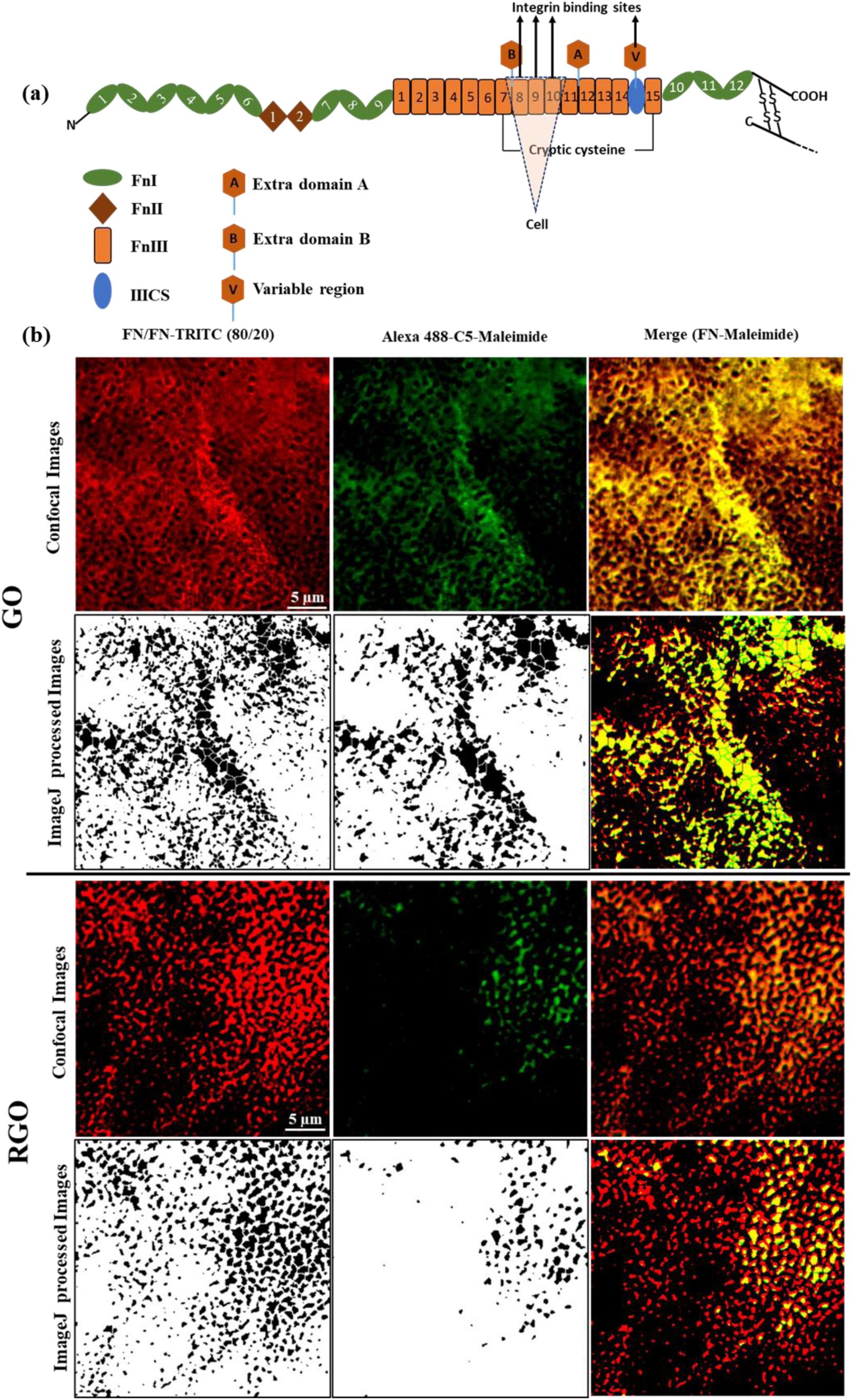
GO shows increased exposure of FN binding sites compared to RGO. (a) Schematic representation of FN molecular structure with different motif domains and (b) Fluorescent micrographs of MA stained FN on GO and RGO surface

The cryptic cysteines on FN are about 3 amino-acid residues from the cell-binding Arg-Gly-Asp (RGD) site that engages cellular integrin receptors.^39^ Formation of a fibrillary FN network plays a critical role in integrin mediated cell adhesion and differentiation. Our results showed that FN on GO provides such FN fibrillar network without any surface modification. In addition to cryptic cysteines, it is possible that the enhanced FN unfolding on GO also exposed hidden cell binding domains (RDG motifs), which may further facilitate better cell attachment and proliferation. To determine if cryptic RGD sites were also exposed by FN adsorption on GO, we developed an assay for integrin attachment to FN-coated GBMs. Prominent RGD-binding integrins (αVβ3 and α5β1) on mesenchymal cells were coated on polystyrene beads and incubated with FN-coated GO and RGO substrates. After extensive washing, we imaged the attached beads on each substrate to measure the relative amount of exposed integrins for each substrate. We found that bead coated with both integrin showed selectively increased attachment for FN-coated GO compared to FN-coated RGO. αVβ3 and α5β1 coated beads showed on average 5- and 3-fold more attachment, respectively, on FN coated GO (**Figure 5**). Thus, the change in graphene chemistry (from GO to RGO) not only affected orientation and molecular conformation of adsorbed protein (FN), but also influenced activity (bioactivity) of protein (FN).

**Figure 5:**
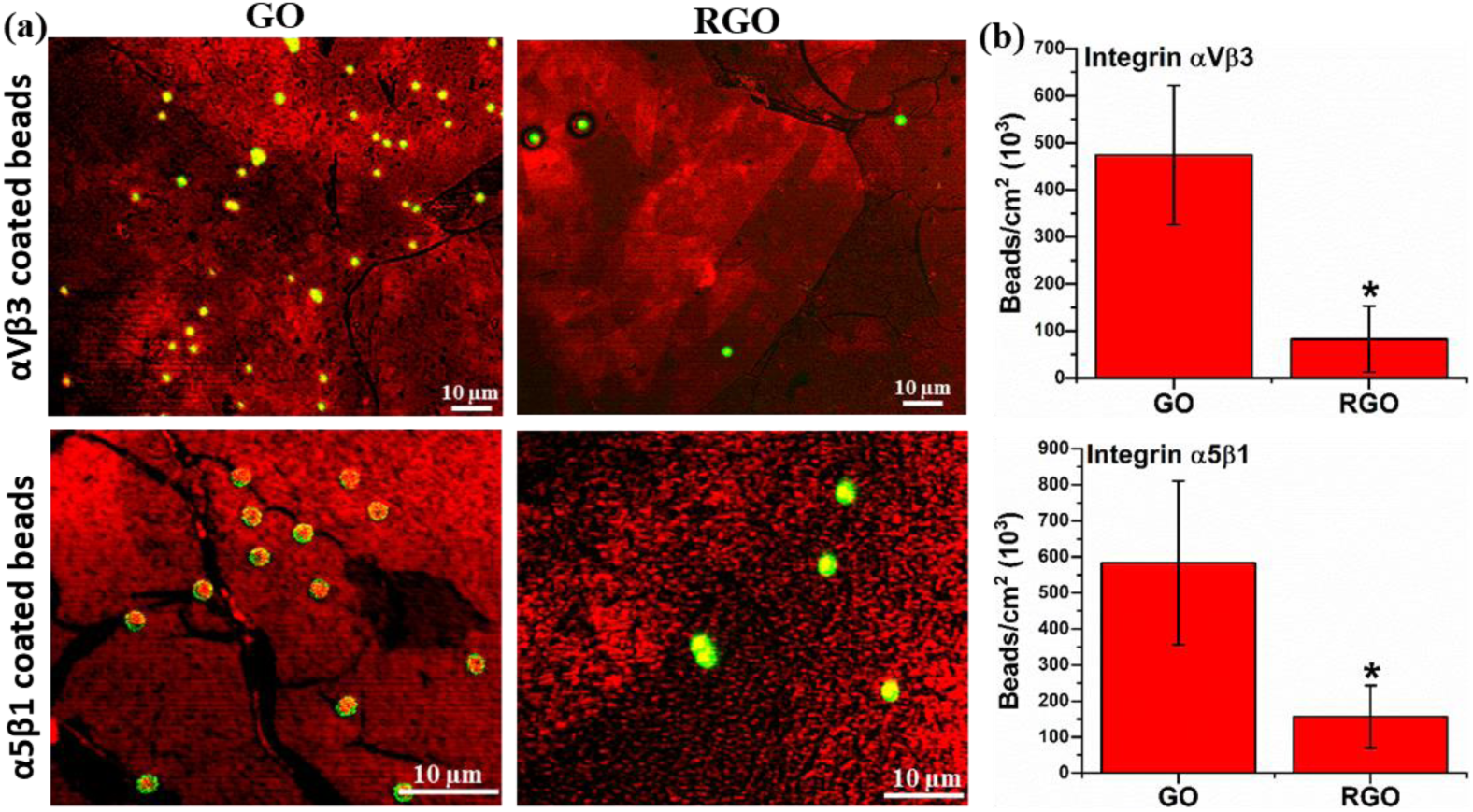
Integrin binding on bioactive RGD site of FN on GBMs. (a) Fluorescence micrograph showing binding of integrin (α_V_β3 and α_5_β1) coated beads on FN adsorbed GO and RGO substrate and (b) Quantification of beads per area on FN adsorbed GO and RGO substrate. Statistically significant differences (p<0.05) compared to GO and RGO are indicated by ‘*’.

### 2.4. Stem cell response on fibronectin coated GO and RGO

Selective binding of both integrins to FN on GO clearly shows that GO interaction with FN not only exposes cryptic cysteines, but also functional RGD motifs that engage integrins. To determine if enhanced integrin binding on GO promoted differential cellular response, we investigated the long-term effect of adsorbed FN on human mesenchymal stem cell (hMSC) attachment, proliferation, and fate. The bioactivity of FN coated GO and RGO with regard to hMSC attachment and proliferation was studied on pristine GO and RGO and FN-coated GO and RGO substrates, under both serum-free (direct) and serum containing (indirect) conditions.

Under direct conditions (with no serum), the presence of FN on GO increased hMSC binding compared to FN-coated on RGO after day 1 of culture (see Supplementary Information for experimental details). hMSCs cultured on all substrates under direct conditions showed minimal increase in DNA amount after day 7, indicating limited proliferation due to serum starvation (**Figure 6(a)**). The absence of serum has been shown to inhibit cell proliferation due to cell cycle arrest and promote senescence.^40^ Overall from day 1 to 7, stem cells on neat GO and FN coated GO showed better attachment and proliferation in comparison to RGO and FN-coated RGO (**Figure 6(a)**). We note that hMSC attachment on neat (-FN-Ser) GO and RGO occurred even in absence of serum (**Figure 6(a)**). Cell attachment on bare substrates are mediated by combinatorial effects of electrostatic, hydrophobic, and non-specific interaction with cell surface molecules.^41^ When coating surfaces with FN (+FN-Ser), it simulated adhesion and influenced cell spreading on GO but not on RGO – where it was inhibitory for cell adhesion. We surmise that the stimulatory effect of adsorbed FN on GO surface arose from elongated fibrillar FN structure with cryptic RGD domain exposed for integrin interaction. As FN on RGO was unfolded to a lesser degree, fewer cryptic RGD domains were exposed decreasing cell attachment.

**Figure 6:**
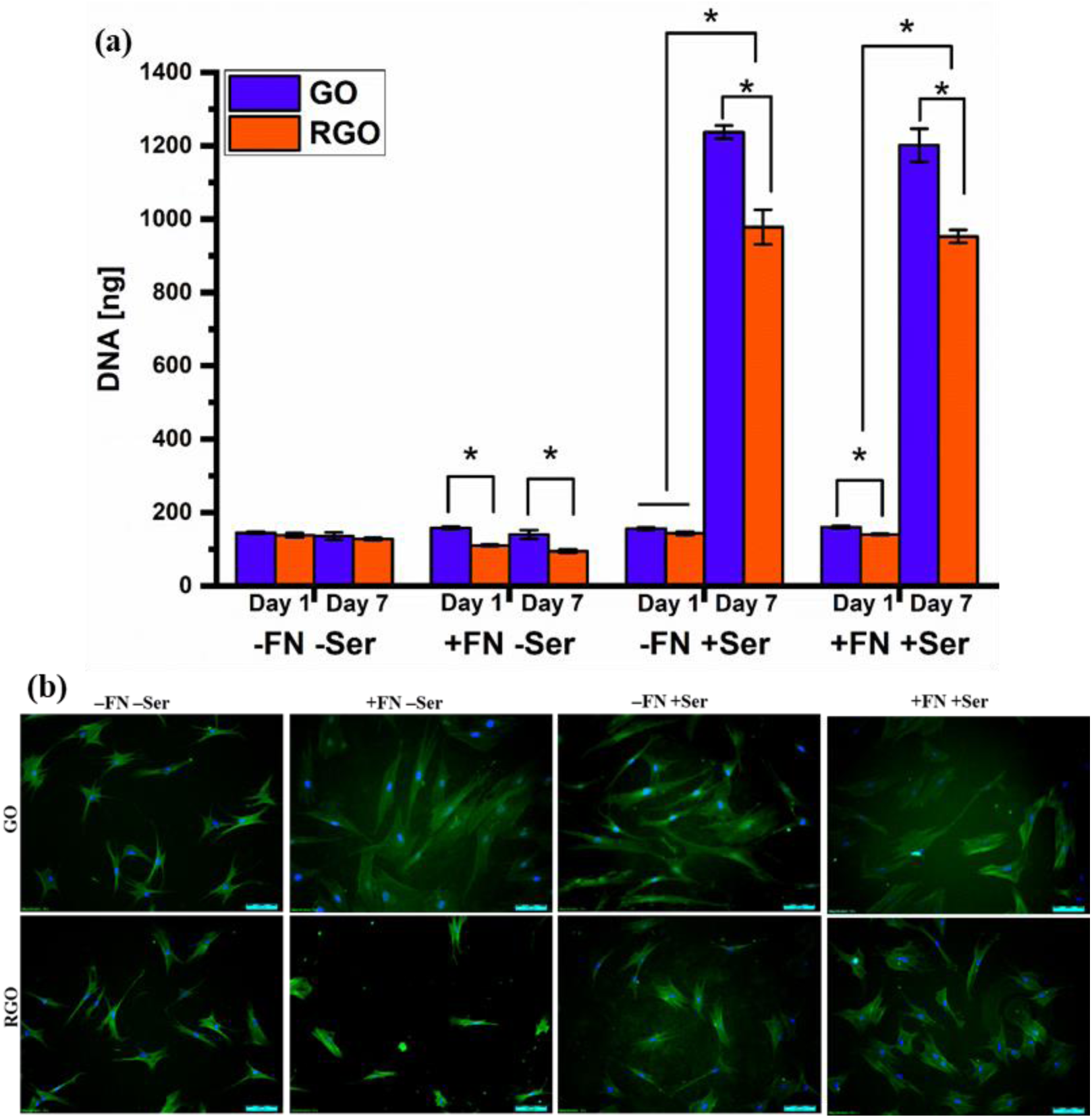
hMSCs attachment, proliferation and morphology on different GBMs surface. (a) Picogreen DNA quantification of hMSCs at day 1 and 7 and (b) Fluroscence staining (Actin and Nucleus) of hMSCs at day 1 (scale bar = 100 µm). Statistically significant differences (p<0.05) compared to GO and RGO are indicated by ‘*’.

We also evaluated the early effects (at day 1) of FN coating GBMs for hMSCs in indirect experiments (in the presence of serum). Surprisingly, in presence of serum, FN-coated RGO (+FN+Ser) and pristine RGO without FN (-FN+Ser) had similar effects on stem cell response in terms of cell attachment and morphology at day 1 (**Figure 6(a) and (b)**). On other hand, FN-coated GO (+FN+Ser) and control GO (-FN+Ser) indirect experiments resulted in a slight change in morphology between the two conditions. hMSCs on FN-coated GO (+FN+Ser) showed 39% larger area in comparison to cells on pristine GO (-FN+Ser) **Figure S5(a)**. In addition, hMSCs on pristine GO (-FN+Ser) showed just 8% more attached cells in comparison to pristine RGO (-FN+Ser), but the morphology of attached cells was significantly different. Cells on pristine GO (-FN+Ser) were highly elongated with an aspect ratio of 5.25, whereas cells on pristine RGO (-FN+Ser) showed less elongation with low aspect ratio of 1.79 (**Figure S5(b)**). In these indirect experiments with serum, the effect of FN coating on hMSCs was quite different for GO and RGO substrates at day 7. We observed more proliferation onto FN-coated GO compared to RGO (**Figure 6(a)**). These results show that addition of FN and serum together provides additional benefit for cell adhesion and growth on GO surfaces. The strong proliferation on both RGO and GO for the indirect experiments can be attributed to additional protein adsorption onto the GBMs (see detail in Supplementary Information; **Figure S6 (a) and (b)**). It has been shown that GO surfaces adsorb more than 50 types of serum protein that can provide a multivalent surface for cell attachment.^42,43^. It is unclear whether the identity of the adsorbed proteins or the strength of adsorption is more important, but our results suggest that both can have strong effects. The significant difference in stem cell response with FN-coating on GO suggests that the pre-adsorbed FN with increased cryptic RGD exposure enhanced hMSC response.

Following testing of FN-coated GBMs with hMSC attachment and proliferation, we tested the ability of FN-coated GBMs (and uncoated GMBs) to elicit a differentiation phenotype. Alkaline phosphatase (ALP), an early marker bone differentiation, showed maximal activity in hMSCs on FN-coated GO in presence of serum. At day 3, hMSCs on pristine GO and RGO (i.e – FN–Ser) and on FN-coated GO and RGO (i.e +FN–Ser) showed negligible ALP activity in absence of serum. On other hand, in presence of serum, hMSCs on pristine GO and RGO (i.e –FN+Ser) showed only faint expression of ALP, which was increased in presence of FN (i.e +FN+Ser) (**Figure 7(a)**). While hMSCs at day 7 showed negligible ALP activity in absence of serum on both pristine (i.e –FN–Ser) and FN-coated (i.e +FN–Ser) GO and RGO substrates, cultures in the presence of serum showed much stronger ALP expression, with FN-coated GO showing the most ALP staining (**Figure 7(b))**.

**Figure 7:**
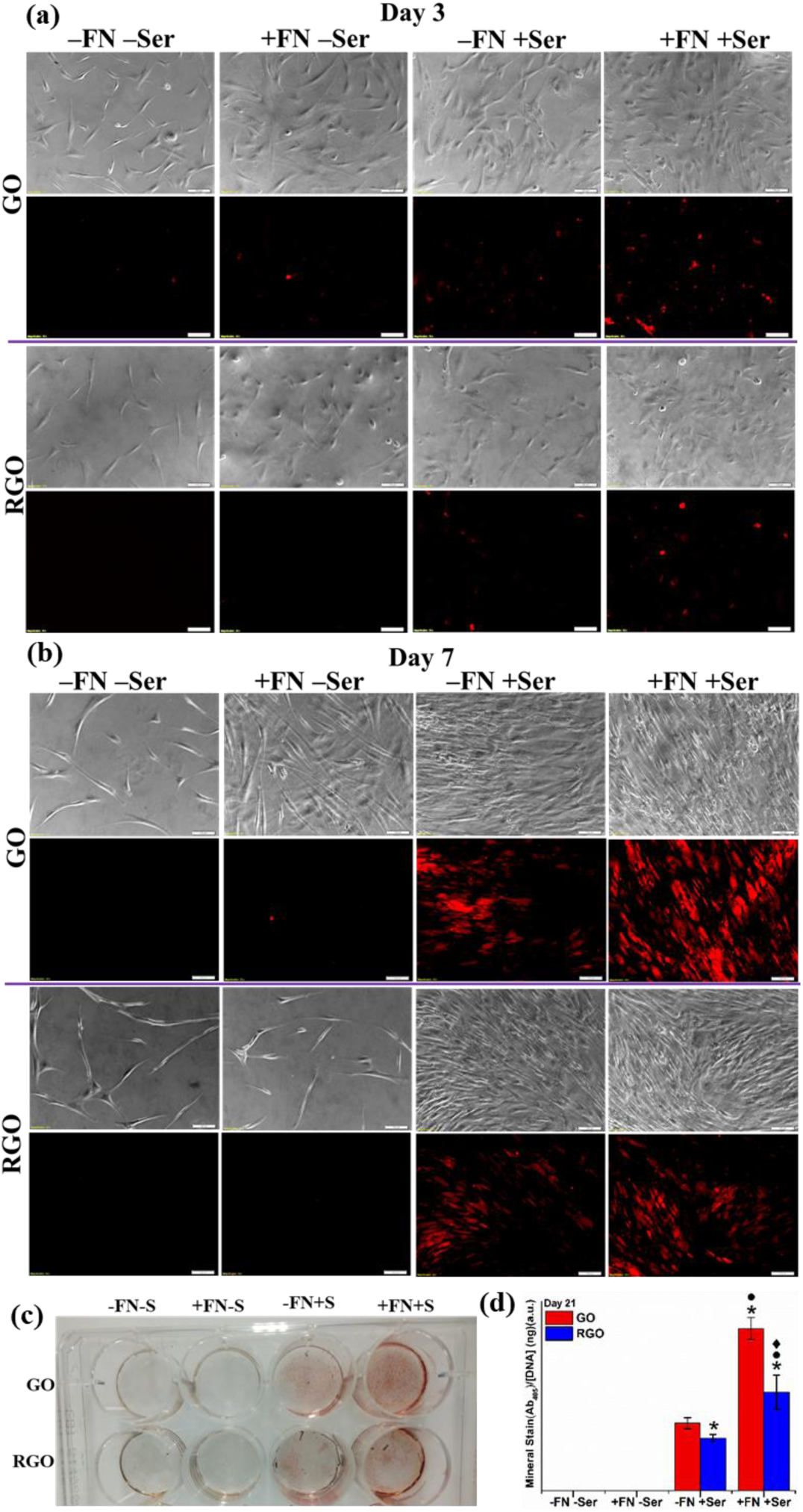
Osteogenic differentiation of hMSc on different GBMs; ALP activity and Mineralization of hMSc. (a) ALP activity at day 3 and (b) at day 7 on neat GO, neat RGO and FN coated GO and RGO substrates in presence and absence of serum (scale bar = 100 µm). (c) digital image of mineral stained by ARS dye and (d) Quantification of calcium deposition by hMSC on neat RGO and FN coated GO and RGO substrates in presence and absence of serum. Statistically significant differences (p<0.05) compared to GO (-FN+Ser), RGO (-FN+Ser), GO (+FN+Ser) and are indicated by ‘*, • and ♦.

While ALP is an early marker for bone differentiation, mineralization is a late stage marker for osteogenesis. We find negligible calcium mineral deposition by hMSCs in absence of serum on all substrates while the presence of serum boosted mineralization significantly on GO compared to RGO (**Figure 7(c)**). In the presence of serum neat GO (-FN+Ser) surface showed around 29% more mineralization compared to RGO (-FN+Ser) whereas after FN coating of GO (+FN+Ser) we found 65% more mineralization in comparison to RGO (+FN+Ser). Thus, FN coating on GO surface improved 140% more mineralization and ALP relative to pristine GO (**Figure 7(d)**). Overall, a small change in GBMs surface chemistry had significant effect on interfacial adsorbed protein from its morphology to conformation, which directly and indirectly influenced stem cell adhesion and fate.

### 2.5. Proposed mechanism for how GBM chemistry affects fibronectin and stem cell fate

Our experiments unequivocally show that FN adsorption is distinct on different GBMs and that is the catalyst for differences in bioactivity and cell response to these materials. Below we describe a mechanism connecting the GMB physicochemistry, protein adsorption and molecular structure and cell fate. During initial adsorption, FN on hydrophilic GO surfaces interacted weakly and formed elongated structures thus undergoing molecular conformational changes, leading to exposure of cryptic bioactive (RGD) sites **(Figure 8(a))**. On other hand, FN on hydrophobic RGO surfaces (with less oxygenated functional groups) interacted strongly forming a compact molecular structure with hidden bioactive sites **(Figure 8(b)).** As a result, stem cell integrins (α5β1 and αVβ3) interacted more on FN coated GO surface resulting in enhanced attachment and cell spreading in absence of serum (a direct interaction indicating how interfacial protein influence cell response). In addition, due to the elongated fibrillar FN structure on GO, it may not only expose RGD sites but also several other multi domains for interaction with other proteins and growth factors present in serum **(Figure 8(c))**. Indeed, pre-coated FN-GO substrates showed strong serum adsorption forming rigid layer of biochemically rich (ECM-like) microenvironment influencing better stem cells differentiation in comparison to FN-RGO **(Figure 8(d))**. Overall, subtle changes in graphene chemistry can have significant impact on interfacial adsorbed protein from its morphology to conformation, which can directly and indirectly influence stem cell fate. Slight changes in GBMs physicochemical properties can have significant effect on interfacial protein, which is the main direct source of interaction with cells on given material.

**Figure 8:**
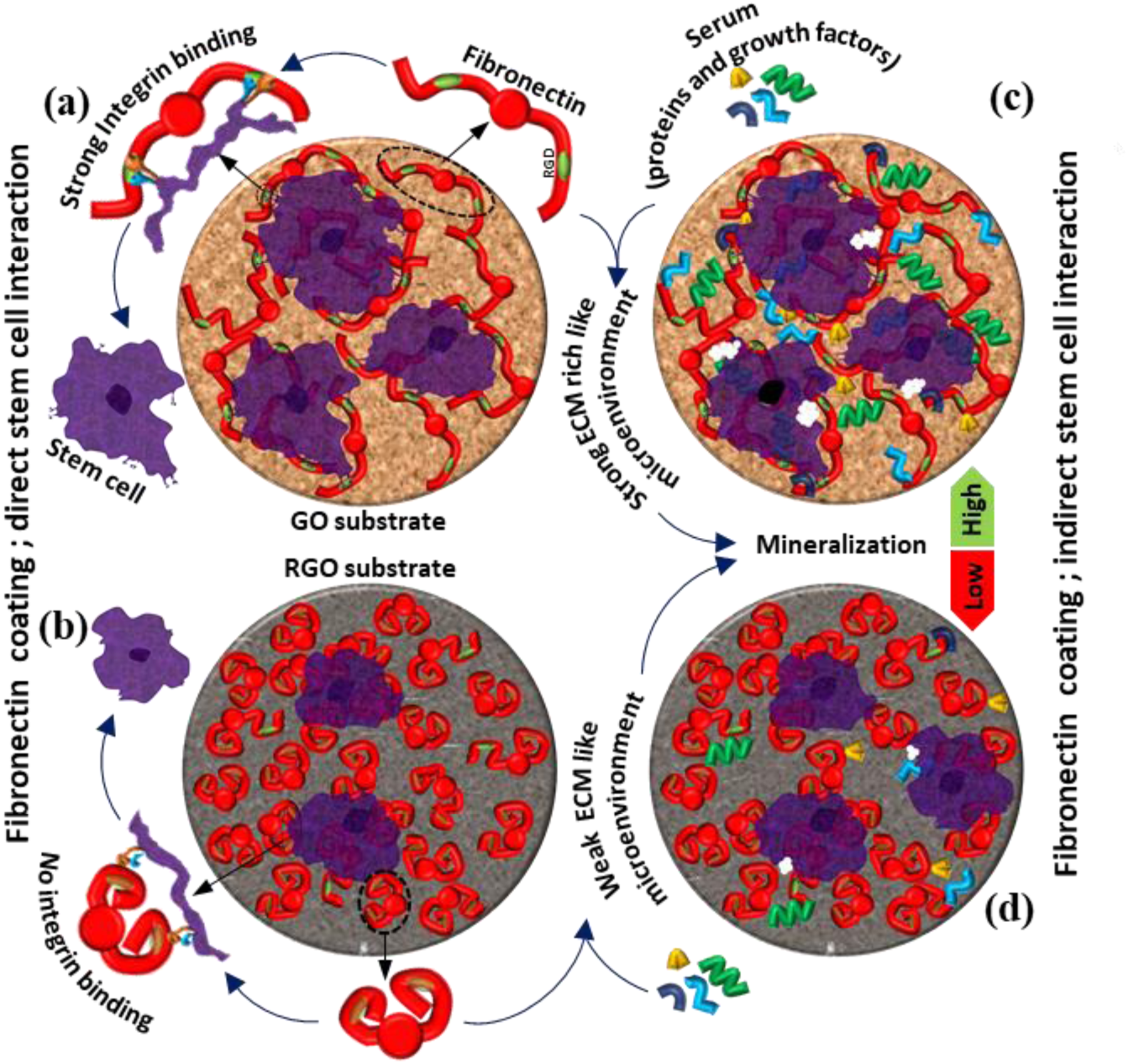
Proposed mechanism for enhanced stem cell ontogenesis on FN coated GBMs. (a) Elongated FN on GO with exposed RGD site for integrin interaction. (b) Compact FN on RGO with cryptic RGD site limiting integrin interaction. (c) Elongated fibrilar FN structure on GO adsorbs more and interact strongly with other proteins and growth factors present in serum forming rigid layer of biochemical rich (ECM like) microenvironment for stem cells differentiation. (d) Compact FN structure on RGO adsorb less and interact weakly with other proteins and growth factors present in serum forming weak layer of biochemical microenvironment for stem cells differentiation.

## 3. Conclusion

GBMs such as GO and RGO – only having different chemical surface properties – showed distinct influence on stem cell fate, as has been previously shown. We further clarified the origin of this effect by quantifying how the physicochemical differences between the GBMs modified adsorption, interaction, and structural conformation of FN from the molecular scale to the macroscale. The hydrophilic and oxygen-rich surface of GO lead to interaction with FN causing formation of elongated fibrillar structures that appeared to form a protein network. On other hand, hydrophobic RGO surfaces (having much fewer oxygenated functional groups) resulted in densely packed and compact FN molecules. Distinct molecular structures of FN on GO and RGO surfaces resulted in different in exposure of integrin binding sites and hMSC interaction. Elongated fibrillar structures of FN on GO showed not only helped hMSC better attachment, but also appeared to promote a rich biochemical microenvironment (by strongly adsorbing serum proteins) that enhanced osteogenic differentiation of hMSCs. Coupling the experiments with biochemical activity and cell response experiments, we demonstrate a multiscale link between physicochemical properties of GBMs, FN unfolding, altered biochemical activity, and stem cell differentiation. Hence, this study highlights how different GBM having physicochemical properties can directly influence protein interaction, which in turn provides a unique biochemical microenvironment to cells for interaction. Tuning GBM physicohemistry is a powerful method to steer cell fate with minimal effort yet exquisite control.

## 4. Experimental Section

### GO and RGO coated glass substrates preparation by spin-coating technique

Graphene Oxide (GO) (a gift from Akimitsu Narita, Max Planck Institute for Polymer Research) was used for all experiments. GO and reduced graphene oxide (RGO) coated glass substrates were prepared by spin coating. For preparation of RGO coated glass substrates, spin coated GO substrates were vacuum-heated at 200 °C for 6 h. (Detail preparation method and schematics provided in Supplementary Information)

### Characterization of prepared GO and RGO coated substrates

Surface coverage of GO and RGO flakes on the glass was characterized by scanning electron microscope (SEM, LEO Gemini 1530, Germany) and atomic force microscopy (AFM, Bruker Dimension Icon) to evaluate surface topography (roughness and thickness) of GO and RGO spin coated sheets. X-ray photoelectron spectroscopy (XPS, Omicron) was used to evaluate different surface chemical functional groups present on prepared GO and thermally treated RGO substrates. Raman spectroscopy was used to evaluate microstructure and structural defect presence on the GO and RGO substrate. Spin coated GO and RGO films were characterized by Raman spectroscopy (uRaman, TechnoSpex) at room temperature with 633 nm laser as an excitation source. Surface wetting properties of spin coated GO and RGO film were evaluated with static water contact angle measurements.

### Fibronectin protein adsorption and characterization

GO and RGO substrates were placed in 12-well plates and Fibronectin (FN) from human plasma (Sigma-Aldrich, 10 µg/ml in PBS) solution was added and allowed to adsorb for 1 hour at room temperature. Coated substrates were gently rinsed with PBS several times to remove weakly adsorbed and non-adsorbed FN prior to any further experiments. Adsorbed FN on GO and RGO substrates was characterized and quantified using XPS. Surface coverage, morphology and structure of adsorbed FN on GO and RGO substrate was studied using AFM in AC mode.

In addition, real-time measurement of FN adsorption on GO and RGO surfaces was performed using quartz crystal microbalance with dissipation monitoring (QCM-D). In a typical experiment, the QCM-D sensor (QSensor QSX 303 SiO_2_) was first cleaned and spin coated with GO and RGO sheets as described above. The prepared QCM-D sensors were flushed with PBS at constant flow rate of 0.1 ml/min for 10 min to establish a baseline. After stabilization of the QCM-D baseline in PBS, the FN solution (10 µg/ml in PBS) was added and adsorption measurement monitored in real time by the change in resonance frequency and dissipation of the QCM sensor due to the adsorbed mass of FN. Finally after rinsing with water, the adsorbed mass density of FN on GO and RGO substrates was calculated in accordance to Sauerbrey equation^44^: Δm = −(C/n)Δf, where C= 17.7 ngcm^−2^s^−1^, n = overtone number and Δf= frequency change. For data analysis, overtone number (n = 3) was used.

### Characterization of Fibronectin morphology and molecular structure

Photo-induced force microscopy, a technique combining nanoscale imaging of AFM with vibrational spectroscopy (Park Scientific) was used to analyze the infrared (IR) spectrum of adsorbed FN on GO and RGO substrates with high spatial resolution. We first scanned the sample in an AC mode in air under ambient conditions to obtain morphology of FN at the nanoscale. Next, Nano-IR images were obtained by scanning the AFM tip across the surface while irradiating the sample at a fixed wavenumber of 1550 cm^−1^, which corresponds to the Amide II band and was selected due to limited laser power at Amide I band (wavenumber of 1650 cm^−1^).

Single spot Nano-IR spectra were collected from 800 to 1800 cm^−1^ with a spectral resolution of 1 cm^−1^ and spatial resolution of 10 nm. Spectra reported in the paper were averaged from at least three measurements from different positions. Nano-IR spectra were normalized against quartz substrate signal, and the amide II band was decomposed using OriginPro software by setting component peak locations found by second-derivative analysis of amide II band. Gaussian functions were used for each sub-peak for spectral decomposition. Peak areas were determined using standard curve-fitting procedures in OriginPro.

### Labelling, visualization, and quantification of Fibronectin cryptic cysteines

Surface exposed cryptic cysteines upon FN adsorption on GO and RGO surface were visualized by incubating the GO or RGO substrates with thiol-reactive maleimide-conjugated Alexa 488 (MA) at 10 µM in PBS. FN-adsorbed GO and RGO substrates were incubated with MA for 1 hour at room temperature. Each substrate was rinsed several times to remove unbound MA before imaging using a laser scanning confocal microscopy (LSM 510, Zeiss). Confocal fluorescence images of adsorbed FN (80:20; FN: rhodamine labelled FN) and bound MA were collected by exiting with laser 550 nm and 488 nm, respectively, using 40X, NA 1.0 water dipping objective. ImageJ was used for image quantification via thresholding of the MA. Confocal fluorescence images were thresholded using the “auto” method. On binarized images, watershed segmentation was performed, and the total segmented area for FN and bound MA were measured. The ratio of bound MA to FN segmented area was expressed as percent labeling of MA on FN-adsorbed GO and RGO surfaces.

### Cell studies

Primary bone-marrow derived human mesenchymal stem cells (hMSCs) (Lonza) were cultured in alpha minimum essential media (alpha-MEM, Invitrogen) supplemented with 15% fetal bovine serum (FBS, Gibco), 1% glutamine and 1% antibiotic mixture of penicillin–streptomycin (Sigma). Cells were incubated at 37° C under 5% CO_2_ atmosphere. The culture media was replaced every three days until cells reached ∼70% confluence. For cell passaging and subculture, cells were removed using 0.25% Trypsin (Gibco). Prior to seeding cells, neat GO and RGO coated substrates were placed in 12 well plates and sterilized with 70% ethanol and allowed for drying. FN (20 µg/ml for GO and 5 µg/ml for RGO) was coated as described above. 25000 cells were seeded in presence and absence of serum in each well. For all cell culture studies neat (uncoated), sterilized GO and RGO surface used as controls.

### Cell attachment, morphology and proliferation

hMSC attachment, morphology, and proliferation on different GBMs surface were studied using a combination of DNA quantification (Picogreen, ThermoFisher) and fluorescence imaging. The Picogreen assay was performed in accordance to previous reported literature,^45^ and fluorescence imaging of hMSCs on different GBMs was done on fixed and permeabilized samples using 3.7% formaldehyde and 0.2% TritonX-100 (Sigma), respectively. Cellular actin filaments were stained with Alexa 488 Phalloidin (100 nM) for 30 min followed by nuclear staining using (14.3 mM) DAPI for 5 min, respectively. The stained cells were imaged using an inverted epi-fluorescence microscope (IX-81, Olympus) using a 20X, 0.4 NA objective with cellSens software. ImageJ was used to evaluate cell morphology. Cells were thresholded to create outlines in order to measure cell area and aspect ratio by fitting an ellipse to each cell as reported previously.^46^

### Binding assay of integrin-coated microbeads to GBM-immobilized FN

The integrin binding activity of FN on GO and RGO was tested with an integrin-coated bead assay. Recombinant human α5β1 and αVβ3 integrins (R&D Systems Bio-Tech) (100 µg/ml) and fluorescent microbeads (Fluoresbrite-Carboxylate, 1.75µm diameter) suspended in PBS (10^8^ beads/ml) were mixed after vortexing beads. α5β1 and αVβ3 (20 µg/ml), respectively, was added to suspended beads (10^6^ beads/ml) and placed on a shaker at 300 rpm for 24 hr at 4°C. Beads were then collected by centrifugation (10000 rpm for 5 min) and washed twice with PBS to remove unbound integrin. Integrin-coated beads were blocked with 2.5% BSA-FITC (Sigma) solution for 24 hr at 4°C to limit non-specific adsorption. Finally, integrin-coated beads were re-suspended in PBS (10^6^ beads/ml) having 1 mM MnCl_2_ for each experimental study. For characterization of bound proteins (α5β1 integrin, αVβ3 integrin, and BSA-FITC) to the microbeads. Detailed explanation on characterization of bound proteins to the microbeads is provided in the Supplementary Information.

For the integrin-FN binding assay, FN-coated GO and RGO substrates were placed in 12-well plates and covered with integrin-coated beads, allowed to bind at room temperature for 1 hour followed by rinsing several times with PBS to remove unbound beads. Rinsed samples were imaged using confocal laser scanning microscopy (LSM 510).

### Alkaline Phosphatase (ALP) activity and differentiation

ALP expression of hMSCs on pristine and FN-coated GO and RGO substrates was studied in presence and absence of serum. Pristine and FN-coated GO and RGO substrates were placed in 12-well plates and seeded with hMSCs (1×10^5^ cells per well) in media in the presence and absence of serum. After 3 and 7 days, cells on different GO and RGO substrates were collected and incubated with BioTracker™ CyP AP dye (10 μM) (Millipore) in MEM medium for 30 min at 37 °C followed by PBS washing.^47^ Cellular ALP activity catalyzes cleavage of the phosphate group in CyP, emitting fluorescence in the near-infrared spectrum, which was quantified using laser scanning microscopy (LSM 510) with an excitation wavelength at 635 nm and emission wavelength at 680-780 nm in the spectral detector.

To evaluate the osteogenic differentiation ability of neat GO and RGO, FN-GO and FN-RGO, hMSCs (1×10^5^ cells per well) were cultured on respective surfaces in absence and presence of serum without any osteogenic supplements for 21 days. *In vitro* mineralization was analyzed by quantifying the calcium deposition at 21 day using 2% Alizarin red S (ARS, Sigma) dye as reported in literature.^48^ For quantification, the ARS stain was dissolved in 0.2 mL of 5% SDS in 0.5 N HCL for 30 min and absorbance of the solubilized stain was measured at 405 nm using a plate reader (Tecan, Germany).

### Statistical analysis

Significant differences between the groups were analysed using 1-way ANOVA (analysis of variance) with Tukey’s test for multiple comparisons used for statistical analysis. Differences were considered statistically significant for p < 0.05 and indicated by symbols in the figure.

## Supporting Information

Supporting information (SI) for this article is available

## Supplement data

1. *GO and RGO coated glass substrates preparation by spin-coating technique*
2. *Physicochemical characterization of GO and RGO substrates*
3. *Fibronectin protein adsorption on GO and RGO coated substrate*
4. *Characterization of bound proteins to the microbeads*
5. *Stem cell response on FN coated GO and RGO substrate*
6. *Serum adsorption and interaction with fibronectin coated GO and RGO substrate*

## Supplement Figures and Tables

***Figure S1:*** *Preparation of GO and RGO substrates. (a) Schematic of GO and RGO coating on glass substrates by spin coating. (b) Digital image of neat UVO treated glass, GO coated glass after spin coating, and RGO coated glass after thermal reduction. (c) Optical micrograph of outlined area (in ‘b’) for neat UVO treated glass, GO coated and RGO coated glass substrate.*

***Table S1***: *Carbon and Oxygen element composition of GO and RGO substrate*

***Figure S2:*** *Physical and chemical characterization of GBMs. (a) AFM profile of surface coverage thickness of GO on glass substrate. (b) SEM micrographs of GO and RGO on glass show flaky coverage. (c) AFM images show surface morphology of GO and RGO flakes on glass. (d) XPS spectra showing chemical functional groups present on prepared GO and thermally treated RGO substrates. (e) Raman spectra showing G and D bands for GO and RGO sheets.*

***Figure S3:*** *Fibronectin adsorption on GBMs. (a) XPS spectra of FN adsorption on GO and RGO surface. (b) AFM images of FN coverage with respective optimized concentration on GO*

***Figure S4***: *Visualization of bound proteins on micro-beads. (a) Digital image of SDS page of integrin coated micro-beads and (b) Fluorescence micrograph of integrin coated FTIC-BSA blocked beads*

***Figure S5***: *Morphology of stem cell on different GBMs surface. (a) and (b) Cell areas and Aspect ratio of hMSCs on neat GO, RGO and FN coated GO and RGO substrates in Presence and absence of serum.*

***Figure S6***: *Serum adsorption and sequential interaction. (a) Adsorption of serum and (b) sequential FN and serum adsorption QCM profile on GO and RGO surface.*

## Acknowledgements

S.K. acknowledges Alexander von Humboldt Foundation Postdoctoral Fellowship, and S.H.P. acknowledges support from the Welch Foundation (F-2008-20190330) and Human Frontiers in Science Program (RGP0045/2018). The authors wish to thank Dr. Rüdiger Berger (MPIP) for AFM imaging, G.Glaßer (MPIP) for SEM imaging and Dr. Volker Mailänder (MPIP) for QCM-D analysis. We thank Anika Keswani for helping to edit this paper.

## Conflict of Interest

The authors declare no competing financial interest.

## References

(1) Goenka, S.; Sant, V.; Sant, S. Graphene-based nanomaterials for drug delivery and tissue engineering. Journal of Controlled Release 2014, 173, 75–88.

(2) Kumar, S.; Chatterjee, K. Comprehensive review on the use of graphene-based substrates for regenerative medicine and biomedical devices. ACS applied materials & interfaces 2016, 8, 26431–26457.

(3) Shin, S. R.; Li, Y.-C.; Jang, H. L.; Khoshakhlagh, P.; Akbari, M.; Nasajpour, A.; Zhang, Y. S.; Tamayol, A.; Khademhosseini, A. Graphene-based materials for tissue engineering. Advanced drug delivery reviews 2016, 105, 255–274.

(4) Kenry, L. W.; Loh, K. P.; Lim, C. T. When stem cells meet graphene: opportunities and challenges in regenerative medicine. Biomaterials 2018, 155, 236–250.

(5) Shi, X.; Chang, H.; Chen, S.; Lai, C.; Khademhosseini, A.; Wu, H. Regulating cellular behavior on few-layer reduced graphene oxide films with well-controlled reduction states. Advanced Functional Materials 2012, 22, 751–759.

(6) Talukdar, Y.; Rashkow, J. T.; Lalwani, G.; Kanakia, S.; Sitharaman, B. The effects of graphene nanostructures on mesenchymal stem cells. Biomaterials 2014, 35, 4863–4877.

(7) Wilson, C. J.; Clegg, R. E.; Leavesley, D. I.; Pearcy, M. J. Mediation of biomaterial–cell interactions by adsorbed proteins: a review. Tissue engineering 2005, 11, 1–18.

(8) Zhang, X.-Q.; Xu, X.; Bertrand, N.; Pridgen, E.; Swami, A.; Farokhzad, O. C. Interactions of nanomaterials and biological systems: Implications to personalized nanomedicine. Advanced drug delivery reviews 2012, 64, 1363–1384.

(9) Li, D.; Zhang, W.; Yu, X.; Wang, Z.; Su, Z.; Wei, G. When biomolecules meet graphene: From molecular level interactions to material design and applications. Nanoscale 2016, 8, 19491–19509.

(10) Kumar, S.; Parekh, S. H. Linking graphene-based material physicochemical properties with molecular adsorption, structure and cell fate. Communications Chemistry 2020, 3, 1–11.

(11) Chong, Y.; Ge, C.; Yang, Z.; Garate, J. A.; Gu, Z.; Weber, J. K.; Liu, J.; Zhou, R. Reduced cytotoxicity of graphene nanosheets mediated by blood-protein coating. ACS nano 2015, 9, 5713–5724.

(12) Guilak, F.; Cohen, D. M.; Estes, B. T.; Gimble, J. M.; Liedtke, W.; Chen, C. S. Control of stem cell fate by physical interactions with the extracellular matrix. Cell stem cell 2009, 5, 17–26.

(13) Lee, W. C.; Lim, C. H. Y.; Shi, H.; Tang, L. A.; Wang, Y.; Lim, C. T.; Loh, K. P. Origin of enhanced stem cell growth and differentiation on graphene and graphene oxide. ACS nano 2011, 5, 7334–7341.

(14) Girase, B.; Shah, J. S.; Misra, R. D. K. Cellular mechanics of modulated osteoblasts functions in graphene oxide reinforced elastomers. Advanced Engineering Materials 2012, 14, B101–B111.

(15) Chaudhuri, P. K.; Loh, K. P.; Lim, C. T. Selective accelerated proliferation of malignant breast cancer cells on planar graphene oxide films. Acs Nano 2016, 10, 3424–3434.

(16) Becerril, H. A.; Mao, J.; Liu, Z.; Stoltenberg, R. M.; Bao, Z.; Chen, Y. Evaluation of solution-processed reduced graphene oxide films as transparent conductors. ACS nano 2008, 2, 463–470.

(17) Eda, G.; Chhowalla, M. Chemically derived graphene oxide: towards large-area thin-film electronics and optoelectronics. Advanced materials 2010, 22, 2392–2415.

(18) Dong, L.; Yang, J.; Chhowalla, M.; Loh, K. P. Synthesis and reduction of large sized graphene oxide sheets. Chemical Society Reviews 2017, 46, 7306–7316.

(19) Slobodian, O. M.; Lytvyn, P. M.; Nikolenko, A. S.; Naseka, V. M.; Khyzhun, O. Y.; Vasin, A. V.; Sevostianov, S. V.; Nazarov, A. N. Low-Temperature Reduction of Graphene Oxide: Electrical Conductance and Scanning Kelvin Probe Force Microscopy. Nanoscale research letters 2018, 13, 139.

(20) Solís-Fernández, P.; Paredes, J.; Villar-Rodil, S.; Martínez-Alonso, A.; Tascón, J. Determining the thickness of chemically modified graphenes by scanning probe microscopy. Carbon 2010, 48, 2657–2660.

(21) Akhavan, O. The effect of heat treatment on formation of graphene thin films from graphene oxide nanosheets. Carbon 2010, 48, 509–519.

(22) Höök, F.; Kasemo, B.; Nylander, T.; Fant, C.; Sott, K.; Elwing, H. Variations in coupled water, viscoelastic properties, and film thickness of a Mefp-1 protein film during adsorption and cross-linking: a quartz crystal microbalance with dissipation monitoring, ellipsometry, and surface plasmon resonance study. Analytical chemistry 2001, 73, 5796–5804.

(23) Liu, S. X.; Kim, J.-T. Application of Kevin—Voigt model in quantifying whey protein adsorption on polyethersulfone using QCM-D. JALA: Journal of the Association for Laboratory Automation 2009, 14, 213–220.

(24) Molino, P. J.; Higgins, M. J.; Innis, P. C.; Kapsa, R. M.; Wallace, G. G. Fibronectin and bovine serum albumin adsorption and conformational dynamics on inherently conducting polymers: a QCM-D study. Langmuir 2012, 28, 8433–8445.

(25) Kennedy, S. B.; Washburn, N. R.; Simon Jr, C. G.; Amis, E. J. Combinatorial screen of the effect of surface energy on fibronectin-mediated osteoblast adhesion, spreading and proliferation. Biomaterials 2006, 27, 3817–3824.

(26) Sengupta, B.; Gregory, W. E.; Zhu, J.; Dasetty, S.; Karakaya, M.; Brown, J. M.; Rao, A. M.; Barrows, J. K.; Sarupria, S.; Podila, R. Influence of carbon nanomaterial defects on the formation of protein corona. RSC Advances 2015, 5, 82395–82402.

(27) Walton, A.; Koltisko, B. Protein-structure and the kinetics of interaction with surfaces. Advances in chemistry series 1982, 245–264.

(28) Silva-Bermudez, P.; Rodil, S. An overview of protein adsorption on metal oxide coatings for biomedical implants. Surface and Coatings Technology 2013, 233, 147–158.

(29) Tellechea, E.; Johannsmann, D.; Steinmetz, N. F.; Richter, R. P.; Reviakine, I. Model-independent analysis of QCM data on colloidal particle adsorption. Langmuir 2009, 25, 5177–5184.

(30) Olsson, A. L.; Quevedo, I. R.; He, D.; Basnet, M.; Tufenkji, N. Using the quartz crystal microbalance with dissipation monitoring to evaluate the size of nanoparticles deposited on surfaces. ACS nano 2013, 7, 7833–7843.

(31) Dixon, M. C. Quartz crystal microbalance with dissipation monitoring: enabling real-time characterization of biological materials and their interactions. Journal of biomolecular techniques: JBT 2008, 19, 151.

(32) Tagaya, M. In situ QCM-D study of nano-bio interfaces with enhanced biocompatibility. Polymer Journal 2015, 47, 599.

(33) Erickson, H. P.; Carrell, N.; McDonagh, J. Fibronectin molecule visualized in electron microscopy: a long, thin, flexible strand. J Cell Biol 1981, 91, 673–678.

(34) Kulik, A. J.; Ruggeri, F. S.; Gruszecki, W. I.; Dietler, G. Nanoscale infrared spectroscopy of light harvesting proteins, amyloid structures and collagen fibres. Microsc Anal 2014, 28, 11–15.

(35) Lindon, J. C.; Tranter, G. E.; Koppenaal, D. Encyclopedia of spectroscopy and spectrometry. (Academic Press, 2016).

(36) Osterlund, E.; Eronen, I.; Osterlund, K.; Vuento, M. Secondary structure of human plasma fibronectin: conformational change induced by calf alveolar heparan sulfates. Biochemistry 1985, 24, 2661–2667.

(37) Thyparambil, A. A.; Wei, Y.; Latour, R. A. Experimental characterization of adsorbed protein orientation, conformation, and bioactivity. Biointerphases 2015, 10, 019002.

(38) Klotzsch, E.; Smith, M. L.; Kubow, K. E.; Muntwyler, S.; Little, W. C.; Beyeler, F.; Gourdon, D.; Nelson, B. J.; Vogel, V. Fibronectin forms the most extensible biological fibers displaying switchable force-exposed cryptic binding sites. Proceedings of the National Academy of Sciences 2009, 106, 18267–18272.

(39) Miyamoto, Y.; Yokoya, A.; Ishizaka, S. Interaction between cell-binding domain and extracellular matrix-binding domain of fibronectin determined by fluorescence depolarization. Biochimica et Biophysica Acta (BBA)-Protein Structure and Molecular Enzymology 1988, 953, 306–313.

(40) Li, B.; Sun, C.; Sun, J.; Yang, M.-h.; Zuo, R.; Liu, C.; Lan, W.-r.; Liu, M.-h.; Huang, B.; Zhou, Y. Autophagy mediates serum starvation-induced quiescence in nucleus pulposus stem cells by the regulation of P27. Stem cell research & therapy 2019, 10, 118.

(41) Hoshiba, T.; Yoshikawa, C.; Sakakibara, K. Characterization of initial cell adhesion on charged polymer substrates in serum-containing and serum-free media. Langmuir 2018, 34, 4043–4051.

(42) Sopotnik, M.; Leonardi, A.; Križaj, I.; Dušak, P.; Makovec, D.; Mesarič, T.; Ulrih, N. P.; Junkar, I.; Sepčić, K.; Drobne, D. Comparative study of serum protein binding to three different carbon-based nanomaterials. Carbon 2015, 95, 560–572.

(43) Qi, Y.; Chen, W.; Liu, F.; Liu, J.; Zhang, T.; Chen, W. Aggregation morphology is a key factor determining protein adsorption on graphene oxide and reduced graphene oxide nanomaterials. Environmental Science: Nano 2019, 6, 1303–1309.

(44) Sauerbrey, G. Use of quartz crystals for weighing thin layers and for weighing. Journal of Physics 1959, 155, 206–222.

(45) Kumar, S.; Raj, S.; Sarkar, K.; Chatterjee, K. Engineering a multi-biofunctional composite using poly (ethylenimine) decorated graphene oxide for bone tissue regeneration. Nanoscale 2016, 8, 6820–6836.

(46) Kumar, S.; Raj, S.; Kolanthai, E.; Sood, A.; Sampath, S.; Chatterjee, K. Chemical functionalization of graphene to augment stem cell osteogenesis and inhibit biofilm formation on polymer composites for orthopedic applications. ACS applied materials & interfaces 2015, 7, 3237–3252.

(47) Li, S.-J.; Li, C.-Y.; Li, Y.-F.; Fei, J.; Wu, P.; Yang, B.; Ou-Yang, J.; Nie, S.-X. Facile and sensitive near-infrared fluorescence probe for the detection of endogenous alkaline phosphatase activity in vivo. Analytical chemistry 2017, 89, 6854–6860.

(48) Kumar, S.; Raj, S.; Jain, S.; Chatterjee, K. Multifunctional biodegradable polymer nanocomposite incorporating graphene-silver hybrid for biomedical applications. Materials & Design 2016, 108, 319–332.

