## Supplemental information for "Molecular control of interfacial protein structure on graphene-based substrates steers cell fate"

#### This supplement information file includes

#### Supplement data

1. *GO and RGO coated glass substrates preparation by spin-coating technique*
2. *Physicochemical characterization of GO and RGO substrates*
3. *Fibronectin protein adsorption on GO and RGO coated substrate*
4. *Characterization of bound proteins to the microbeads*
5. *Stem cell response on FN coated GO and RGO substrate*
6. *Serum adsorption and interaction with fibronectin coated GO and RGO substrate*

#### Supplement Figures and Tables

***Figure S1:*** *Preparation of GO and RGO substrates. (a) Schematic of GO and RGO coating on glass substrates by spin coating. (b) Digital image of neat UVO treated glass, GO coated* *glass after spin coating, and RGO coated glass after thermal reduction. (c) Optical micrograph of outlined area (in 'b') for neat UVO treated glass, GO coated and RGO coated glass substrate.*

***Table S1**: Carbon and Oxygen element composition of GO and RGO substrate*

***Figure S2:** Physical and chemical characterization of GBMs. (a) AFM profile of surface coverage thickness of GO on glass substrate. (b) SEM micrographs of GO and RGO on glass show flaky coverage. (c) AFM images show surface morphology of GO and RGO flakes on glass. (d) XPS spectra showing chemical functional groups present on prepared GO and thermally treated RGO substrates. (e) Raman spectra showing G and D bands for GO and RGO sheets.*

***Figure S3:** Fibronectin adsorption on GBMs. (a) XPS spectra of FN adsorption on GO and RGO surface. (b) AFM images of FN coverage with respective optimized concentration on GO*

***Figure S4**: Visualization of bound proteins on micro-beads. (a) Digital image of SDS page of integrin coated micro-beads and (b) Fluorescence micrograph of integrin coated FTIC-BSA blocked beads*

***Figure S5**: Morphology of stem cell on different GBMs surface. (a) and (b) Cell areas and Aspect ratio of hMSCs on neat GO, RGO and FN coated GO and RGO substrates in Presence and absence of serum.*

***Figure S6**: Serum adsorption and sequential interaction. (a) Adsorption of serum and (b) sequential FN and serum adsorption QCM profile on GO and RGO surface.*

### 1. GO and RGO coated glass substrates preparation by spin-coating technique

GO and reduced graphene oxide (RGO) coated glass substrates were prepared by spin coating. Glass substrates (diameter 22 mm; thickness 0.3 mm (NeoLab)) were cleaned by immersing in a 10% surfactant (Micro90) solution and sonicated for 2 h. Thereafter, the glass was washed thoroughly in ethanol and deionized water, followed by nitrogen drying, and finally treated in an ultraviolet ozone (UVO) chamber for 30 min to render the surface hydrophilic. 200 µl of GO solution (1 mg/ml in $H_2O$) was added to the UVO treated glass and spin coated at 2000 rpm for 30 sec with 200 rpm/sec acceleration. Samples were then dried under vacuum for 24 h at room temperature. For Preparation of RGO coated glass substrates, spin coated GO substrates were vacuum-heated at 200 °C for 6 h. A schematic illustration for the preparation of spin coated GO and RGO film substrate is shown in **Figure S1(a)**.

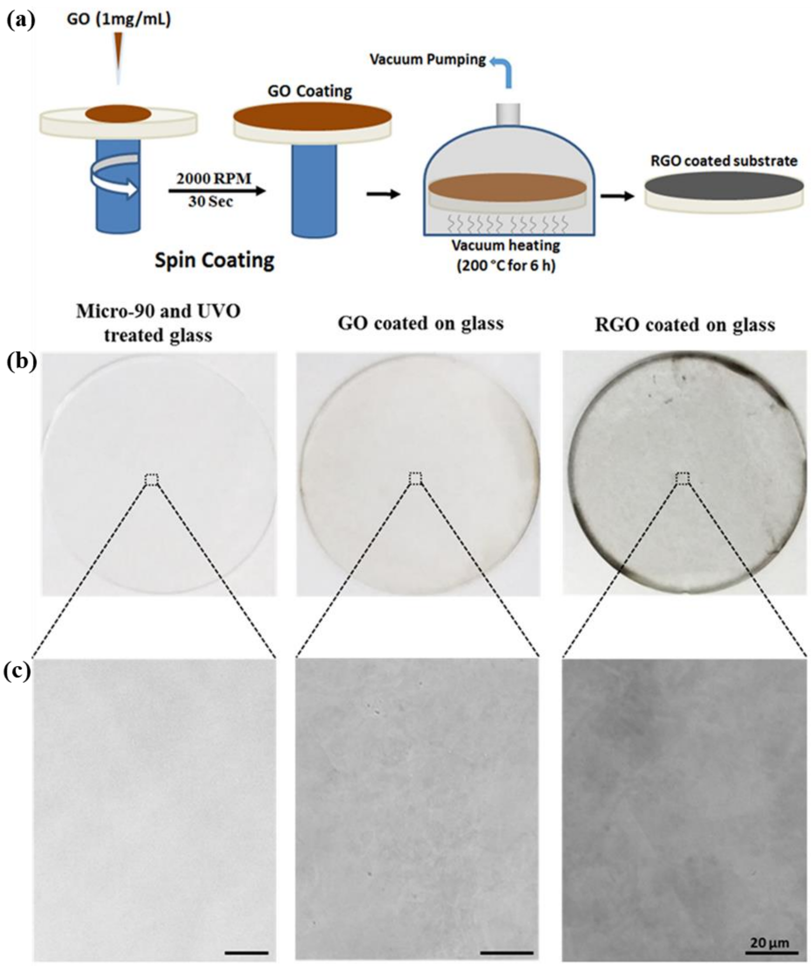


**Figure S1:** Preparation of GO and RGO substrates. (a) Schematic of GO and RGO coating on glass substrates by spin coating. (b) Digital image of neat UVO treated glass, GO coated glass after spin coating, and RGO coated glass after thermal reduction. (c) Optical micrograph of outlined area (in ‘b’) for neat UVO treated glass, GO coated and RGO coated glass substrate.

GO and RGO coated glass substrates were prepared by spin coating GO on to cleaned glass substrates. Prior to spin coating GO, glass substrates were cleaned and UVO treated to enhance surface wettability for strong adhesion of GO sheets on hydrophilic surface.[1,2] **Figure S(b)** shows a digital photograph of UVO-treated, GO coated and RGO coated glass. After coating GO on cleaned UVO treated glass, a semi-transparent brownish color can be seen due to covering of GO sheets. After thermal reduction, the color changed from brownish to black – a visible characteristic of GO reduction. The visible color change from brownish to black for RGO film can be attributed to increased light absorbance of RGO sheets in the visible and near infrared region, likely due to recovery of $sp^2$ hybridization of graphene. [3] Optical micrographs showed nearly transparent flaky sheet coverage of GO while thermally reduced RGO sheets showed much more opaque contrast (**Figure S(c)).**

**2. Physicochemical characterization of GO and RGO substrates**

SEM showed coverage of GO and RGO sheets on glass substrates (**Figure S2(b)**). GO flakes lay flat on a glass surface without wrinkles. After thermal treatment, RGO flakes did not lead to any obvious morphological changes and maintained a compact, wrinke-free structure (**Figure S2(b) inset**). AFM of the spin coated GO and RGO sheets on UVO treated glass surface are shown in **Figure S2(c)**. These images show the irregular sheets of GO and RGO with lateral dimensions ranging from a few to several micrometers with a surface coverage thickness of ~ 5.5 nm (**Figure S2(a)**). This shows that GO and RGO coating contained approximately 3-5 monolayers of stacked sheets. Both GO and RGO coating showed flakes that were smooth and continuous with distinguishable edges of individual sheets. The average surface roughness ($R_a$) of GO and RGO coated substrate was 0.6 nm and 0.4 nm, respectively, suggesting that RGO film was slight smoother than GO potentially due to annealing after thermal treatment. [4] Importantly we found that thermal reduction of GO to RGO did not produce any significant morphological difference apart from apparent decrease in flakes thickness, which is consistent with previous work showing that the thickness of an individual GO sheet reduced from 1.1 nm to 0.8 nm after thermal reduction.[5,6]

**Figure S2(d)** shows the characteristic XPS spectra of GO and RGO with C1s and O1s peaks at 285 and 535 eV, respectively. After thermal treatment of GO the relative peak intensity ratio of C1s/O1s increased, corresponding to an increase in atomic content of carbon to oxygen due to removal of oxygenated functional groups in reduction. The (C/O) ratio increased from 1.19 to 2.51 for RGO upon thermal removal of oxygenated functional groups as shown in **Table S1**.

| | **Carbon (At %)** | **Oxygen (At %)** | **C/O (%)** |
|---|---|---|---|
| **GO** | 54.4±8.7 | 45.6±8.4 | 1.19 |
| **RGO** | 71.9±2.6 | 28.1±2.5 | 2.51 |

**Table S1**: Carbon and Oxygen element composition of GO and RGO substrate

Further analysis of C1s core-level spectra of GO and RGO by spectral decompostion into various chemical states using a Gaussian fitting after base line correction[7] shows the change in core C1s spectra shapes before and after thermal treatment, supporting the notion that reduction leads to a change in chemical composition. Overall, these results indicate that a large fraction of oxygenated functional groups from GO surface was removed upon thermal treatment.

**Figure S2(e)** shows Raman spectra of GO and RGO with respective existence of D and G bands. GO showed presence of G band at 1590 $cm^{-1}$ whereas RGO showed slight shift in G band to 1586 $cm^{-1}$. The shift in G band for RGO is an indicative of reduction of GO during thermal annealing.[8] Furthermore both GO and RGO showed D band at 1325 $cm^{-1}$, indicative of the presence of defects or disorder in the graphene materials. [9,10] The presence of oxygenated functional groups on basal plane and at edges of graphene sheets creates structural defects by distorting translational and periodic symmetry of graphene $sp^2$ carbon network.[11,12] Measuring $I_D/I_G$ ratio is indicative of presence of defects in graphene sheet. $I_D/I_G$ ratio decreased from 1.5 (for GO) to 1.3 (for RGO) after thermal reduction, in alignment with our XPS results.

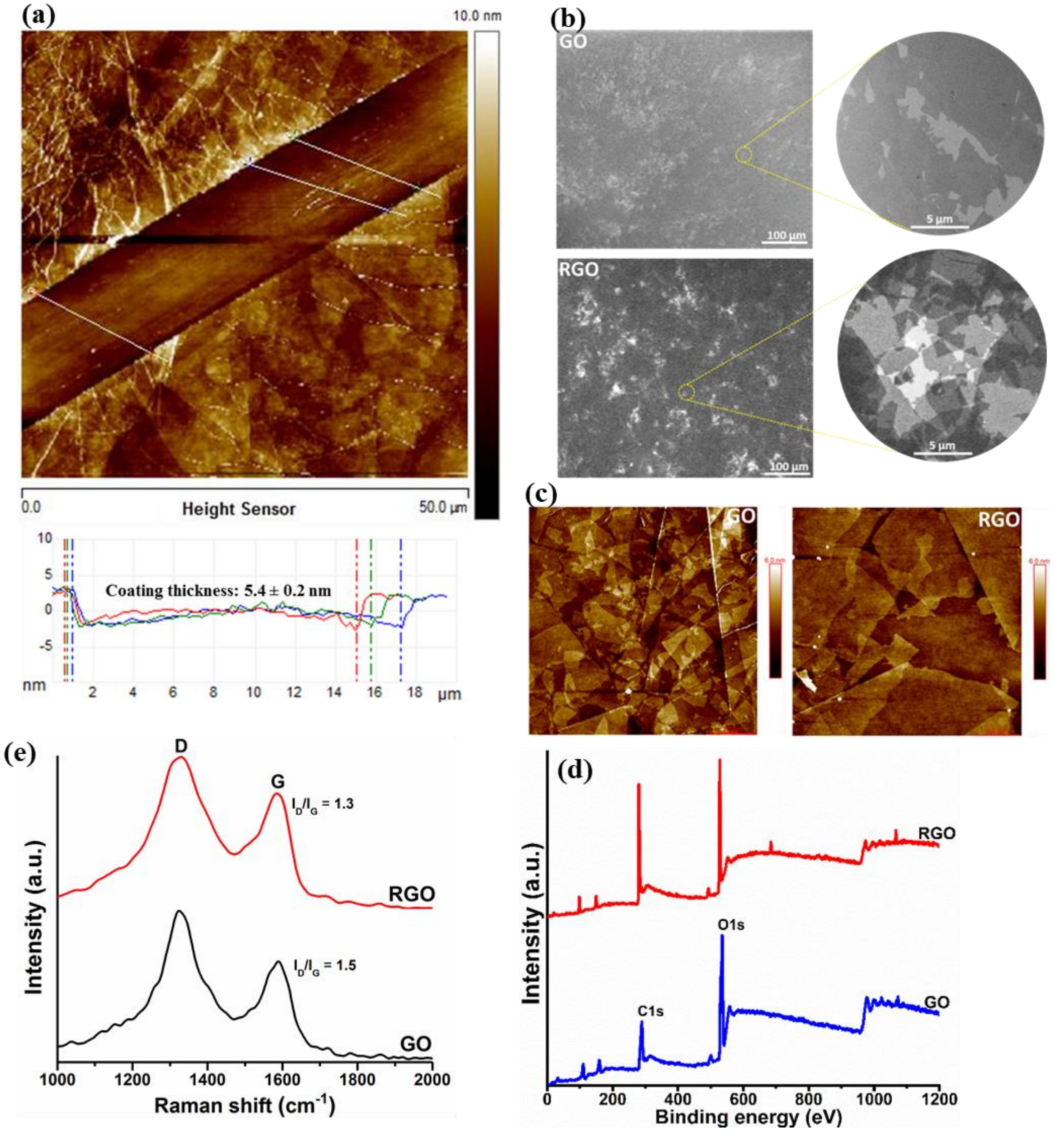


**Figure S2:** Physical and chemical characterization of GBMs. (a) AFM profile of surface coverage thickness of GO on glass substrate. (b) SEM micrographs of GO and RGO on glass show flaky coverage. (c) AFM images show surface morphology of GO and RGO flakes on glass. (d) XPS spectra showing chemical functional groups present on prepared GO and thermally treated RGO substrates. (e) Raman spectra showing G and D bands for GO and RGO sheets.

### 3. Fibronectin protein adsorption on GO and RGO coated substrate

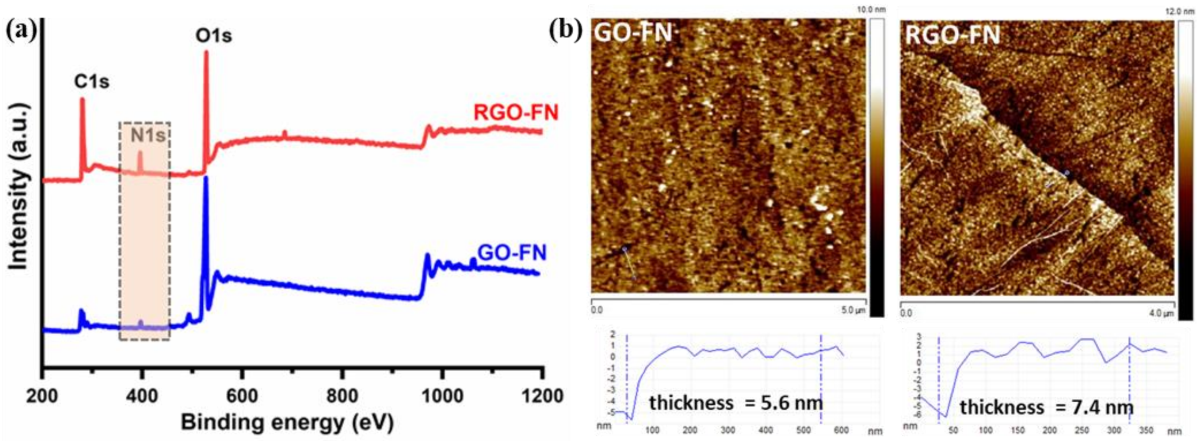


**Figure S3:** Fibronectin adsorption on GBMs. (a) XPS spectra of FN adsorption on GO and RGO surface. (b) AFM images of FN coverage with respective optimized concentration on GO (20 µg/ml) and RGO (5 µg/ml) surface.

**Figure S3(a)** shows XPS spectra of FN adsorbed onto the GO and RGO substrates. Of immediate interest in these spectra is the presence of nitrogen (N 1s) elemental peak, which can only come from proteins in our measurements. The N 1s peak has been routinely used for protein quantification in XPS measurements.[13] RGO showed relatively intense peak for nitrogen with 6.9% atomic weight percentage in comparison to GO having 1.4% atomic percentage of nitrogen. This clearly indicates a nearly fivefold more FN adsorption on RGO with respect to GO. After optimization of FN concentration to achieve homogenous adsorption. AFM FN adsorption profile on GO and RGO showed uniform coverage and thickness of 5.6 nm and 7.4 nm for GO and RGO, respectively.

### 4. Characterization of bound proteins to the microbeads

Proteins bound (integrin α5β1, αVβ3 and BSA-FITC) microbeads were run in a SDS gel by immersing them in gel loading buffer (NuPAGE LDS) under reducing conditions. The beads in the sample buffer were loaded on 4–12% gradient gels (NuPAGE) and run against pure integrin α5β1, αVβ3 and BSA-FITC as controls, along with high-molecular-weight marker (ranging from 10 to 225 kDa (Novagen)), as described previously[14]. In addition, laser scanning confocal microscopy (LSM 510) was used to visualize BSA-FITC coated on fluorescent microbeads.

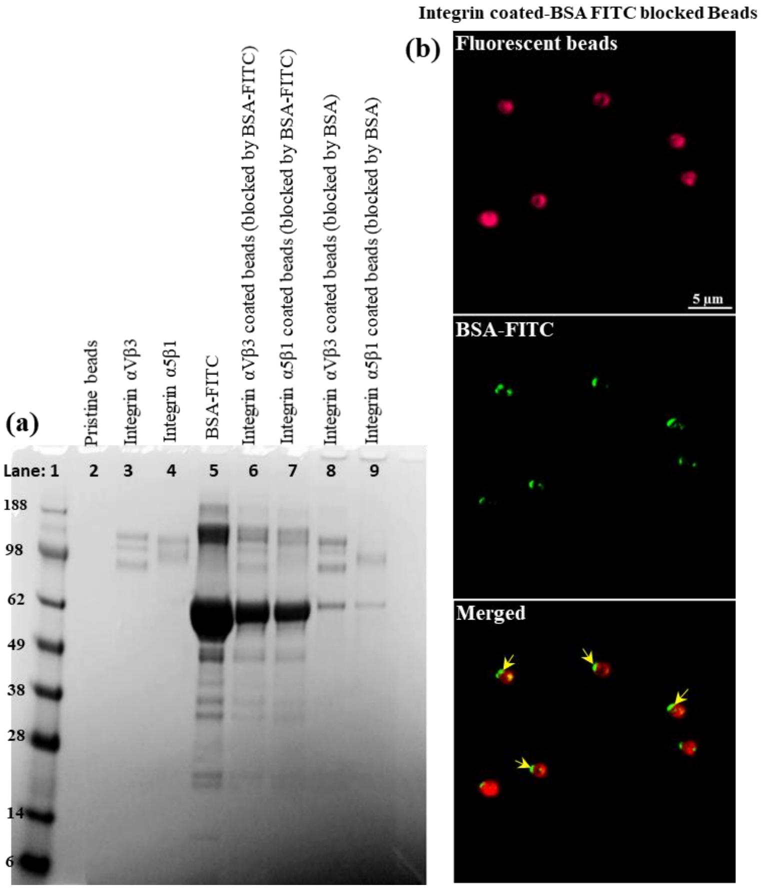


**Figure S4**: Visualization of bound proteins on micro-beads. (a) Digital image of SDS page of integrin coated micro-beads and (b) Fluorescence micrograph of integrin coated FTIC-BSA blocked beads

Bound proteins on microbeads were characterized using SDS gel and fluorescence imaging. **Figure S4(a)** shows SDS-PAGE pattern of standard protein marker, pristine beads, integrin αVβ3, α5β1, BSA-FITC as control and integrin (αVβ3 and α5β1) coated beads with BSA-FITC and normal BSA blocking (in sequence from lane 1 to 9). Lane 1 shows standard protein bands for different molecular weight markers ranging around 188 kDa to 6 kDa. Lane 2 with pristine beads showed presence of no bands suggesting no traces of any protein impurities present in purchased fluorescent microbeads. Lane 3 showed characteristic protein bands for control integrin αVβ3 at around 130-100 kDa for $\alpha_V$ subunit and β3 subunit. Similarly lane 4 also showed 2 protein bands adjacent to each other at around 120-95 kDa for $\alpha_5$ subunit and β3 subunit. The protein bands of integrin αVβ3 and α5β1 fall in range of prescribed molecular mass of respective (α) and (β) subunits described by supplier (R&D System). Lane 5 showed presence of protein bands form commercial BSA-FITC; interestingly, BSA-FITC showed multiple bands with intense broad band around 62 kDa. Multiple SDS page bands for BSA FITC can be attributed to the multimer nature of BSA and/or presence of impurities in commercial BSA-FITC as reported by several researchers.[15,16] Lane 5 to 9 revealed presence of protein bands at similar position to that of control integrin (lane 3 for αVβ3 and lane 4 for α5β1) along with BSA-FITC and control BSA bands. These results clearly indicated coating of integrin αVβ3 and α5β1 on micro beads along with BSA blocking. In addition, confocal laser scanning micrograph for BSA-FITC blocked integrin coated beads showed fluorescent beads (red) having green spots for BSA-FITC protein (indicated by arrow)(**Figure S4(b)**) on the periphery signifying nonspecific binding of BSA on beads during blocking.

**5. Stem cell response on fibronectin coated GO and RGO substrate**

Under direct conditions (in absence of serum) after 1 day of culture, stem cells on the control (-FN-Ser) GO and RGO (-FN-Ser) surface were comparable in number (based on DNA amount) with similar morphology. On the other hand hMSCs on FN-coated (+FN-Ser) GO and RGO showed differences in cell number and morphology. hMSCs on FN-coated GO showed ~40% more attached cells with a spread, elongated morphology having a larger area in comparison to hMSCs on FN coated RGO (**Figure S5(a)and (b)**). Furthermore, in comparison to neat (-FN-Ser) RGO, FN coated (+FN-Ser) RGO decreased cell adhesion by 26% whereas FN coated (+ FN-Ser) GO showed 13% cell attachment compared to neat (-FN-Ser) GO. These results suggest that the presence of FN on GO (in the absence of serum) increases interaction with hMSCs resulting in more cell attachment.

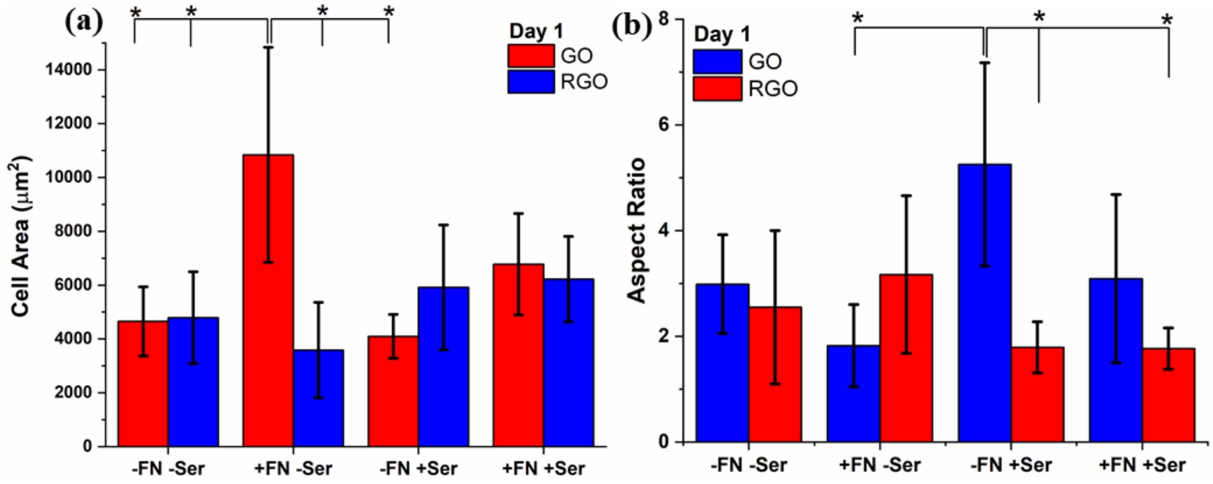


**Figure S5**: Morphology of stem cell on different GBMs surface. (a) and (b) Cell areas and Aspect ratio of hMSCs on neat GO, RGO and FN coated GO and RGO substrates in Presence and absence of serum.

We also evaluated the early effects (at day 1) of FN coating GBMs for hMSCs in the presence of serum. Stem cells on control (-FN+Ser) GO surface showed just 8% more attached cells in comparison to (-FN+Ser) RGO, but the morphology of attached cells was significantly different. Cells on (-FN+Ser) GO were highly elongated with an aspect ratio of 5.25, whereas cells on (-FN+Ser) RGO showed less elongation with low aspect ratio of 1.79 (**Figure S5(b)**).

FN coated GO (+FN+Ser) substrates in presence of serum showed an increase (15% more) in cell attachment in comparison to FN coated RGO (+FN+Ser). In terms of cell morphology, cells on FN-coated GO were slight larger compared to cells on FN-coated RGO surface (both in the presence of serum). Surprisingly, in presence of serum, FN-coated RGO (+FN+Ser) and control RGO (-FN+Ser) surface had similar effects on stem cell response in terms of cell attachment and morphology at day 1. This is in contrast to FN-coated GO (+FN+Ser) and control GO (-FN+Ser) in presence of serum where we observed a slight change in morphology. hMSCs on FN-coated GO (+FN+Ser) showed 39% larger area in comparison to cells on neat GO (-FN+Ser) **Figure S5(a)-(b)**. These results show that addition of FN and serum together provides additional benefit for cell adhesion on GO surfaces.

**6. Serum adsorption and interaction with fibronectin coated GO and RGO substrate**

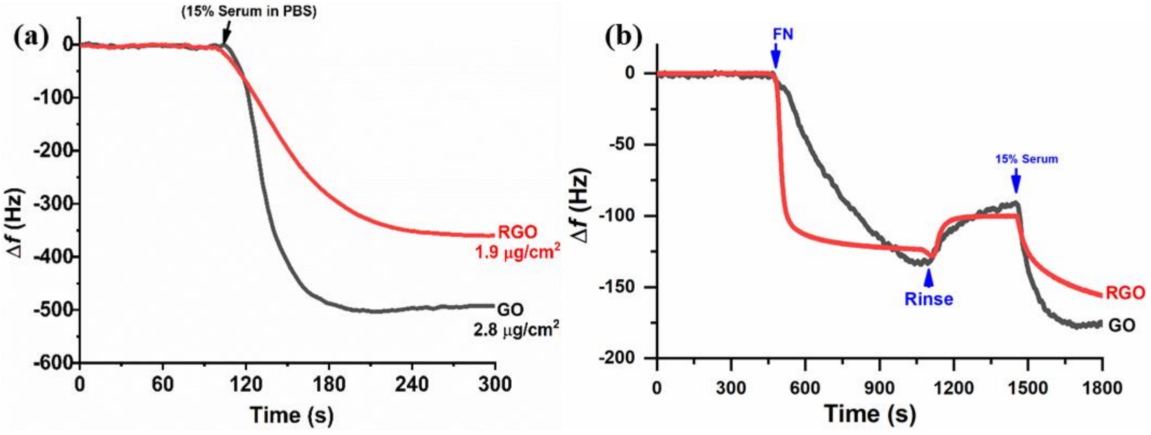

**Figure S6**: Serum adsorption and sequential interaction. (a) Adsorption of serum and (b) sequential FN and serum adsorption QCM profile on GO and RGO surface.

In order to confirm conformational effect of FN on GO and RGO surface influencing serum protein adsorption and interaction, First we studied adsorption profile of serum on neat GO and RGO surface. **Figure S6(a)** shows fast and high adsorption of serum proteins (2.8 μg/cm$^2$) on GO surface in comparison to (1.9 μg/cm$^2$) on RGO surface. We further evaluated sequential adsorption profile of FN followed by serum on GO and RGO surface **(Figure S6(b))**. Real time FN adsorption profile was similar as described previsouly suggesting almost uniform and complete coverage of FN. Later sequential addition of serum showed fast adsorption of serum components on FN-GO surface in comparison to FN-RGO surface. GO-FN surface showed 489.9 ng/cm2 serum component adsorption in comparison to 312.7 ng/cm2. More serum protein coverage on FN-GO surface clearly distinguish the availability and distribution of serum components to cells. Other interesting observation was that the rapid adsorption profile of serum on FN-GO surface resembled similar to FN adsorption profile on RGO surface, which suggests that serum protein not only adsorb faster but also form rigid layer on FN-GO surface may be due to stronger protein-protein interaction. Similarly, neat GO adsorption profile of serum proteins resembles to that of FN-GO serum adsorption profile suggesting protein-protein interaction mediating fast and strong adsorption.  Formation of rigid layer due to stronger protein-protein interaction can be attributed to extended fibrilar FN structure. These results provide useful insights regarding the probable mechanisms of how rigid ECM based rich biochemical microenvironment influence stem cells fate.

### References for Supplementary Information

(1) Kuang, P.; Constant, K.Increased wettability and surface free energy of polyurethane by ultraviolet ozone treatment. *Wetting and Wettability, M. Aliofkhazraei, ed., InTech, Rijeka, Croatia* **2015**, 85-103.

(2) Robinson, J. T.; Zalalutdinov, M.; Baldwin, J. W.; Snow, E. S.; Wei, Z.; Sheehan, P.; Houston, B. H.Wafer-scale reduced graphene oxide films for nanomechanical devices. *Nano letters* **2008**, 8, 3441-3445.

(3) Robinson, J. T.; Tabakman, S. M.; Liang, Y.; Wang, H.; Sanchez Casalongue, H.; Vinh, D.; Dai, H.Ultrasmall reduced graphene oxide with high near-infrared absorbance for photothermal therapy. *Journal of the American Chemical Society* **2011**, 133, 6825-6831.

(4) Pang, S.; Tsao, H. N.; Feng, X.; Müllen, K.Patterned graphene electrodes from solution-processed graphite oxide films for organic field-effect transistors. *Advanced Materials* **2009**, 21, 3488-3491.

(5) Solís-Fernández, P.; Paredes, J.; Villar-Rodil, S.; Martínez-Alonso, A.; Tascón, J.Determining the thickness of chemically modified graphenes by scanning probe microscopy. *Carbon* **2010**, 48, 2657-2660.

(6) Akhavan, O.The effect of heat treatment on formation of graphene thin films from graphene oxide nanosheets. *Carbon* **2010**, 48, 509-519.

(7) Yang, D.; Velamakanni, A.; Bozoklu, G.; Park, S.; Stoller, M.; Piner, R. D.; Stankovich, S.; Jung, I.; Field, D. A.; Ventrice Jr, C. A.Chemical analysis of graphene oxide films after heat and chemical treatments by X-ray photoelectron and Micro-Raman spectroscopy. *Carbon* **2009**, 47, 145-152.

(8) Chen, W.; Yan, L.Preparation of graphene by a low-temperature thermal reduction at atmosphere pressure. *Nanoscale* **2010**, 2, 559-563.

(9) Kudin, K. N.; Ozbas, B.; Schniepp, H. C.; Prud'Homme, R. K.; Aksay, I. A.; Car, R.Raman spectra of graphite oxide and functionalized graphene sheets. *Nano letters* **2008**, 8, 36-41.

(10) Ferrari, A. C.; Robertson, J.Interpretation of Raman spectra of disordered and amorphous carbon. *Physical review B* **2000**, 61, 14095.

(11) Gupta, B.; Kumar, N.; Panda, K.; Kanan, V.; Joshi, S.; Visoly-Fisher, I.Role of oxygen functional groups in reduced graphene oxide for lubrication. *Scientific reports* **2017**, 7, 45030.

(12) Dreyer, D. R.; Todd, A. D.; Bielawski, C. W.Harnessing the chemistry of graphene oxide. *Chemical Society Reviews* **2014**, 43, 5288-5301.

(13) Ray, S.; Shard, A. G.Quantitative analysis of adsorbed proteins by X-ray photoelectron spectroscopy. *Analytical chemistry* **2011**, 83, 8659-8666.

(14) Rastian, Z.; Pütz, S.; Wang, Y.; Kumar, S.; Fleissner, F.; Weidner, T.; Parekh, S. H.Type I collagen from Jellyfish Catostylus mosaicus for biomaterial applications. *ACS Biomaterials Science & Engineering* **2018**, 4, 2115-2125.

(15) Determan, A. S.; Trewyn, B. G.; Lin, V. S.-Y.; Nilsen-Hamilton, M.; Narasimhan, B.Encapsulation, stabilization, and release of BSA-FITC from polyanhydride microspheres. *Journal of controlled release* **2004**, 100, 97-109.

(16) McDonagh, P. F.; Williams, S. K.The preparation and use of fluorescent-protein conjugates for microvascular research. *Microvascular research* **1984**, 27, 14-27.
